# Synthetic periphyton as a model system to understand species dynamics in complex microbial freshwater communities

**DOI:** 10.1101/2021.10.31.466637

**Authors:** Olga Lamprecht, Bettina Wagner, Nicolas Derlon, Ahmed Tlili

**Affiliations:** Eawag: Swiss Federal Institute of Aquatic Science and Technology, Dübendorf, Switzerland

**Keywords:** Freshwater biofilm, periphyton, benthic algae, synthetic microbiology, microbial composition, amplicon sequencing, multiple stressors, temperature, pesticides

## Abstract

Phototrophic biofilms, also known as periphyton, are microbial freshwater communities that drive crucial ecological processes in streams and lakes. Gaining a deep mechanistic understanding of the biological processes occurring in natural periphyton remains challenging due to the high complexity and variability of such communities. To address this challenge, we rationally developed a workflow to construct a synthetic community by co-culturing 26 phototrophic species (i.e., diatoms, green algae and cyanobacteria) that were inoculated in a successional sequence to create a periphytic biofilm on glass slides. We show that this community is diverse, stable and highly reproducible in terms of microbial composition, function and 3D spatial structure of the biofilm. We also demonstrate the ability to monitor microbial dynamics at the single species level during periphyton development and how their abundances are impacted by stressors such as increased temperature and a herbicide, singly and in combination. Overall, such a synthetic periphyton, grown under controlled conditions, can be used as a model system for theory testing through targeted manipulation.

## Introduction

Periphyton are benthic freshwater communities of high diversity that are dominated by photoautotrophic microorganisms, such as diatoms, green algae and cyanobacteria [1–3]. Periphyton forms the basis of aquatic food-webs as primary producer, contributes to the release of oxygen and influences the hydrodynamics of the water body [3–5]. Due to its ecological importance in stream functioning, periphyton is widely used as a biological community model to assess effects of biotic and abiotic factors, such as trophic interactions, man-made chemical pollution or eutrophication [6]. However, the majority of existing data on periphyton are descriptive thus far, focusing on integrative endpoints, such as photosynthesis or biomass, that basically consider the community as a black box [7]. Such an integrative approach can be useful because it informs on the general community status. It does not, however, reveal mechanisms that govern the structural dynamics and functional properties of natural communities. This notwithstanding, previous studies have showed that in addition to environmental factors, diatom species interactions and microbial succession in natural periphyton play a major role in shaping the final community composition [8, 9].

Gaining mechanistic understanding of the biological processes in periphyton, such as species interactions or responses to environmental factors, remains challenging due to the complexity of the communities [10]. The varying species composition in natural periphyton where species strongly interact with each other further complicates the evaluation of the outcomes. Every periphyton, and natural microbial community in general, is somewhat different owing to multiple and varying environmental factors [1]. Hence, natural microbial communities will, to a major part, remain undefined, despite the increasing power to annotate species by, for example, next generation sequencing [11]. These limitations have inspired the development of synthetic microbial ecology as a discipline [12, 13]. It relies on the establishment of communities that are artificially created by co-culturing selected microbial genotypes while retaining key functionalities and properties of the more complex natural counterparts. These well-defined and relatively simplified consortia of interacting microorganisms can then be used as a model system for hypothesis testing in order to answer key ecological questions. By knowing the exact microbial composition of such reproducible communities, it is possible to monitor precisely changes of each single species within the community over time.

Synthetic microbial communities have been mainly constructed with bacterial biofilms [14–16] and planktonic freshwater algae [17, 18]. These communities were used to examine, for example, species interactions, biodiversity-productivity relationships and effects of environmental stressors [14, 16, 17, 19, 20]. Despite the long and extensive history of research on periphyton and its ecological importance, very few attempts have been made to design and construct a benthic synthetic community with a focus on diatom species [8, 21]. Here we aimed to develop a workflow by co-culturing multiple freshwater phototrophic species to establish a reproducible, complex and tenable synthetic periphyton, which do not only include diatoms but also green algae and cyanobacteria. We applied a bottom-up approach [22] based on rational design and current knowledge on microbial dynamics during periphyton formation. Relying on several criteria, we first selected twenty-six phototrophic microbial species and organized them into functional groups based on microbial succession during periphyton development. Then we defined the starting inoculum for each group, based on the growth rates of each single species composing the groups. Finally, we experimentally determined the colonization duration for each group. The established workflow of sequential addition of the groups to colonize an artificial substrate, covered with an extracellular polymeric substance (EPS)-producing bacteria, is summarized in Fig. 1. As a case study, we used the established synthetic periphyton to test for the single and combined effects of the herbicide terbuthylazine and temperature change on the community spatial structure, function and microbial composition.

**Fig. 1:**
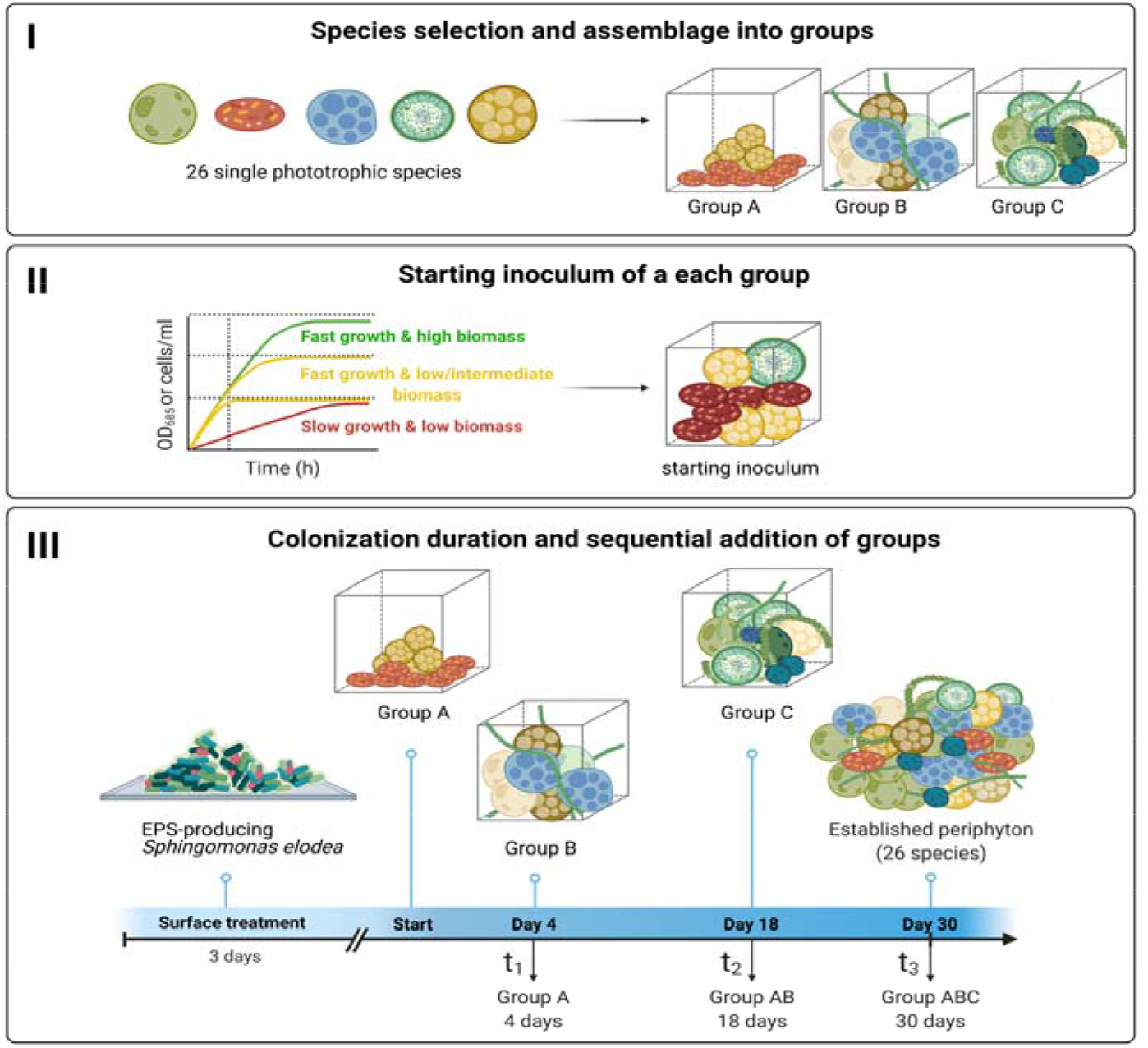
Schematic representation of the workflow to establish the synthetic periphyton. Twenty-six phototrophic microbial species were selected based on their ability to grow under the same conditions and their frequent detection in natural freshwater periphyton. The selected species were then assigned to three groups (**I)**. The cell density in the starting inoculum for each group was determined based on the growth rates and reached maximum biomass of each single species (**II)**. Groups were added sequentially onto growth chambers covered with the EPS-producing bacteria *Sphingomonas elodea*. Time points for the addition of the groups and colonization durations for each phototrophic group were determined experimentally **(III)** (Illustration created with BioRender.com).

## Results and Discussion

### Species selection and grouping criteria

Following a literature search and screening of available databases that describe freshwater periphyton microbial composition, we selected a total of 34 phototrophic species, consisting of 18 diatoms (Bacillariophyta), 10 green algae (Chlorophyta) and 6 Cyanobacteria (Supplementary Table 1). The selection criteria included detection frequency in freshwater periphyton, cultivable species that are commercially available from algal banks, and a taxonomic composition that covers the different stages of periphyton development. Such composition implies various biological traits, such as nutrient requirements and sensitivity to stressors, and consequently different growth conditions. Therefore, we progressively acclimatized all the 34 phototrophic species for at least six months to grow in the same liquid medium (i.e., COMBO, Supplementary Table 2) as well as temperature and light conditions (see Methods for more details). Twenty-six out of the 34 photosynthetic species were successfully adapted and thus selected for the subsequent establishment of the synthetic periphyton (Table 1).

**Table 1.**
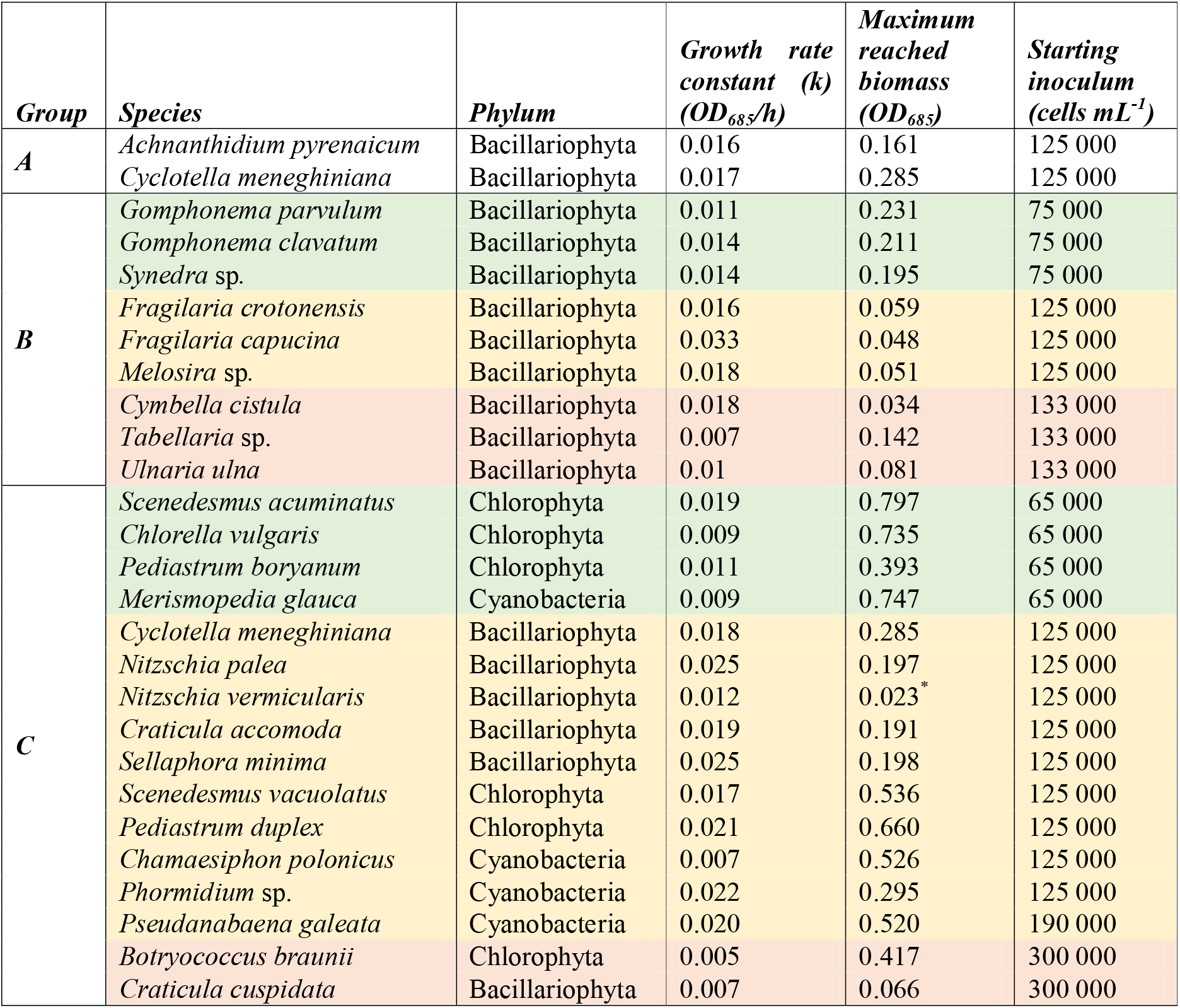
List of the 26 selected phototrophic species used to establish the periphyton. Species were assembled into groups A, B and C, according to the order of addition to the community. Growth rate constant *k* (OD_685_/h) and maximum reached biomass (maximum OD_685_) were derived from optical density measurements at 685 nm (Supplementary Table 3). Color code: green – fast growth and high biomass; yellow – fast growth and low/intermediate biomass; red – slow growth and low biomass. To ensure accurate grouping of species, growth rate constant *k* (cell number/h) and maximum reached biomass (maximum cell number) were additionally derived from CASY cell counter for most of green algae and diatom species (see Supplementary Table 4). *Due to its cell morphology resulting in inaccurate measurements with the spectrophotometer, *N. vermicularis* was included in the intermediate (yellow) group based on the growth rate and maximum biomass measured by CASY cell counter (see Supplementary Table 4).

The initial cell density of single microbial species and their arrival order to a habitat, as well as potential intra- and interspecific competition for light and nutrients, can strongly influence species succession dynamics and community assembly in periphyton [8, 9, 21, 23]. In a closed growth system, such as the one we used in our study (see Method for more details), these processes might lead to a severely reduced diversity within the community. Because one of our main objectives when constructing our synthetic community was to maintain a relatively high level of diversity, we opted for a sequential introduction of the species into the growth chambers. For this, we distributed the twenty-six selected species into three groups (A, B and C) based on the current knowledge about periphyton development and microbial succession dynamics of phototrophs in fresh waters [23–25] (Table 1). Group A consists of *A. pyrenaicum*, a known pioneer diatom in periphyton formation, and *C. meneghiniana*, one of the most commonly found freshwater diatoms. These two species belong to the low profile guild (see also Supplementary Table 1), which comprises the most resistant species with regard to nutrient-poor conditions, and therefore dominate at low nutrient levels [23, 25]. Group B consists of 9 high profile diatoms that contribute the most to the 3D structure development of the periphyton [23]. The high profile guild encompasses species of tall stature, filamentous, chain-forming, stalked diatoms and species with comparatively small cells that form long colonies and extend beyond the boundary layer, i.e. the thin layer of fluid in the immediate vicinity to a periphyton surface under flow conditions. This allows them to exploit resources that are unavailable to the low profile guild [23, 24]. Since our experimental setup was performed under static conditions, the high profile guild diatoms could extend and grow into large structures, possibly only being disturbed by the laminar flow. The remaining 15 species, which compose group C, correspond to the green algae and cyanobacteria as well as the diatoms from the genera *Navicula, Nitzschia* and *Sellaphora* that belong to the motile guild. This guild includes relatively fast moving species, which allows them to physically avoid stress and to move towards resource-rich microhabitats within the benthic mat [24, 26–28]. Green algae and cyanobacteria were added to the periphyton in the last group C, since they are most sensitive to nutrient depletion and light attenuation and might produce unfavorable and harmful toxins to the other added species, respectively [29, 30]. An exception was made for *C. meneghiniana*, which was also included in Group C in addition to group A, as this species is not only low profile but also planktonic and can therefore colonize periphyton at any stage of development. This might explain its high abundance and frequent detection in natural periphyton [23].

Finally, we considered that the first stage of biofilm formation is the attachment of EPS-producing bacteria to the colonization substrate [31, 32]. With the development of genomics techniques, various bacterial strains belonging to the *Sphingomonas* family have been frequently identified as initiators of periphyton formation [33, 34]. We therefore used *Sphingomonas elodea* (ATCC 31461) for surface coating. *S. elodea* is a bacterium known to produce an exopolysaccharide gellan gum with unique colloidal and gelling properties [35]. This bacterium provided a solid and sticky attachment layer for the subsequently introduced phototrophic species, allowing them to bind the surface, multiply and presumably produce their own EPS.

### Starting inoculum and colonization duration for each phototrophic group

As growth rates vary among microorganisms, one major challenge to obtain a complex community is to avoid that one or very few species dominate the community. In order to address this challenge, we defined the starting inoculum for each group by using suspension growth experiments, even though single species growth might differ between planktonic and benthic modes. Therefore, based on the planktonic growth rate and reached planktonic maximum biomass for each single species in suspension (details in Supplementary Tables 3 and 4) we added the least number of cells for species that grew fast and reached high biomasses, and the highest number of cells for species that grew slow and reached low biomasses (as shown in Fig. 1). The total starting cell concentration of all species within a group was 250’000 cells mL^−1^, 999’000 cells mL^−1^ and 2’175’000 cells mL^−1^ for group A, B and C, respectively (see detailed cell concentrations for each species in Table 1).

After determining the starting inoculum, we experimentally tested and set the optimal colonization duration for each of the formed groups. For this purpose, we monitored over time the increase of surface-attached cell number and the PSII quantum yield as proxies for benthic biomass and community function, respectively. As well, we measured surface colonization dynamics based on microscopy imaging. Adequate colonization time was especially important for group A in order to allow the low profile diatoms to colonize the EPS-covered surface but not to overgrowth the colonization substrate. Our results show that when grown separately, *C. meneghiniana* and *A. pyrenaicum* reached similar number of surface-attached cells (Fig. 2A) and the highest PSII quantum yield (Supplementary Fig. 2A) after 4 days of benthic growth. What is more, when both species were grown together at a one to one ratio for 4 days, the large majority of cells (approximately 6 to 7 times more cells than in the supernatant) were attached to the substrate surface (see Fig 2B and 2C). They also covered 9.6 ± 1.4% of the surface (Supplementary Fig. 3), leaving sufficient surface for colonization by the subsequently added groups B and C. Based on these results, we selected 4 days as the optimal colonization duration for group A. However, it is important to mention that in natural periphyton, the colonization substratum might first be entirely covered with the pioneer species, which then are replaced or overgrown by late colonizing species.

**Fig. 2:**
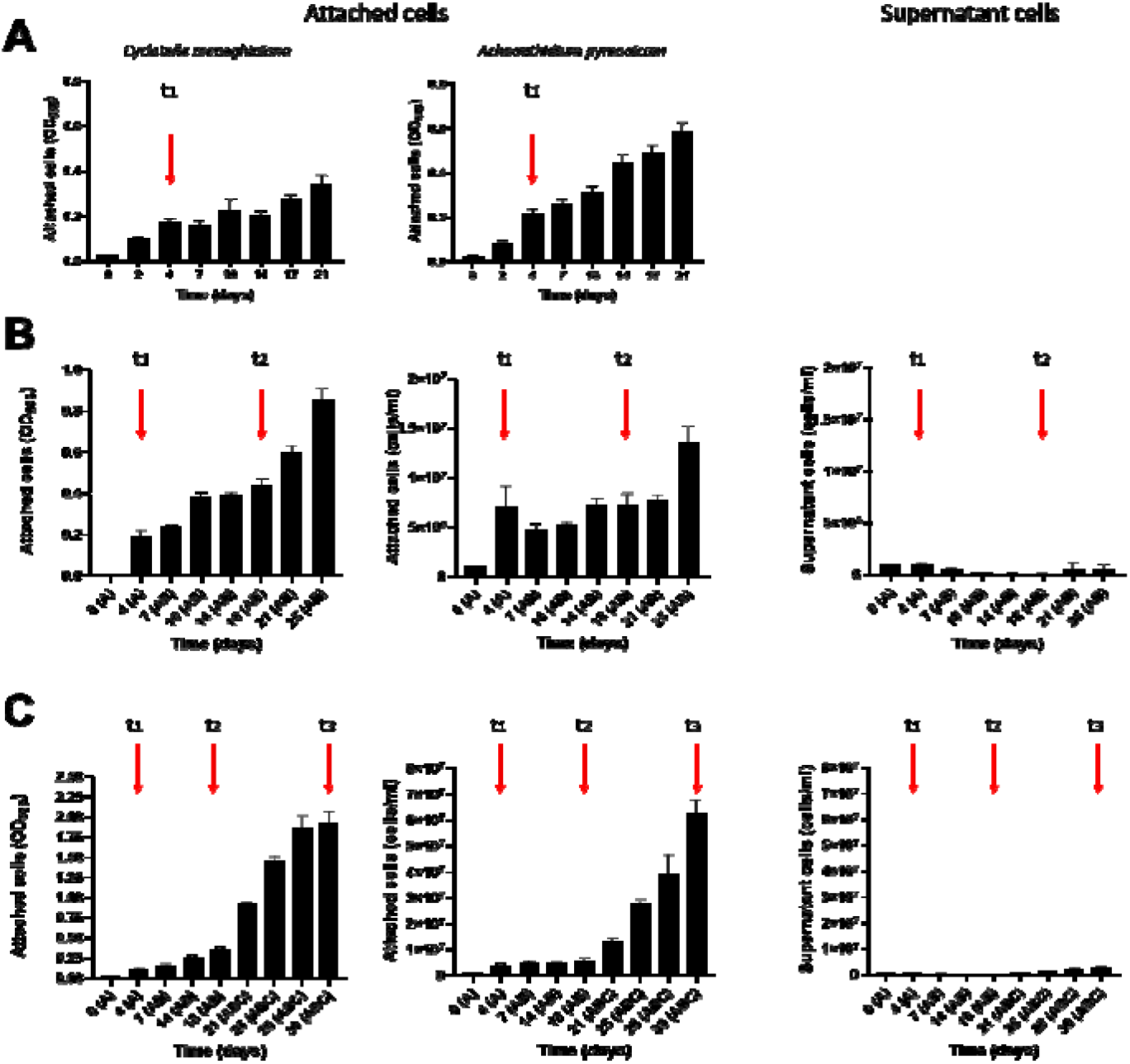
Increase of the benthic biomass during periphyton establishment. The concentrations (means ± SD; n = 3) of attached and non-attached cells (supernatant) were quantified via optical density (OD) at 685 nm and cell count with a CASY cell counter throughout the periphyton development; separately for the two species of group A **(A)**, then for groups A and AB (upon addition of group B onto group A at 4 days of growth) **(B)** and for groups A, AB and ABC (upon addition of group C onto group AB at 18 days of growth) **(C)**. Red vertical arrows indicate selected time points when group B species were added on top of group A (t_1_), group C on top of group AB (t_2_) and periphyton was established (t_3_). Cell concentrations in the supernatant for *C. meneghiniana* and *A. pyrenaicum* when grown separately **(A)** were below the quantification limit; hence no graphics for supernatant is shown.

To determine the colonization duration for groups B and C, we followed a slightly different strategy. Since group B species were added on top of group A after 4 days, we monitored over time the number of cells attached to the substrate and in the supernatant, as well as PSII quantum yield for the group AB together. Our data show that 14 days after group B was added to group A, most of the cells were attached (Fig. 2B) and covered 47.4 ± 3.7 % of the surface (Supplementary Fig. 4). Longer incubation, on the other hand, led to an increase of detached cells (Fig. 2B) and a decrease in the PSII quantum yield (Supplementary Fig. 2B). Such results indicate that the periphyton formed by group AB has reached its maturity and that 14 days is the optimal colonization duration for group B, when added on top of group A. Subsequently, at 18 days (A 4 days + B 14 days) of colonization, we introduced group C. Twelve days after group C was added, most of the cells were attached to the colonization surface (Fig. 2C) with a coverage area of 91.8 ± 1.9 % (Supplementary Fig. 4) and relatively high PSII quantum yield (Supplementary Fig. 2C), indicating that we obtained a stable benthic community after a total colonization duration of 30 days. Such a duration is usually considered optimal for natural periphyton to reach its maturity when growing on artificial and natural substrates [2, 36] and has been applied in most of the field and microcosm studies focusing on periphyton (e.g., [11, 37–39]).

### 3D structural changes during periphyton establishment

The spatial organization of periphyton is of importance as it provides microenvironments and niches for specific microorganisms as well as allows species that are located in close proximity to interact [40]. Here we used optical coherence tomography (OCT), which enables image acquisition at the meso-scale in the range of millimeters (mm) [41] (Fig. 3A) and provides details of overall thickness, surface topology (relative roughness) and internal porosity changes during biofilm formation [42]. At four days (t_1_) of colonization by group A, the periphyton reached an average thickness of 101±47 μm and a relative roughness of 0.4 (Fig. 3B). After species from group B were introduced and grown together with species from group A for another 14 days (t_2_), the average periphyton thickness slightly decreased while the relative roughness and internal porosity increased. This suggests that high profile diatoms (group B) can modify the overall periphyton architecture from a rather homogeneous structure (Ra’ < 0.5) formed by group A species towards a heterogeneous one with a higher surface roughness (Ra’ > 0.5). Finally, following the addition of group C and after 30 days (t_3_) of growth, the established periphyton was almost 200 μm thick and highly dense, as indicated by the sharp thickness increase and internal porosity decrease, respectively. A potential explanation of the increased structure density could be the presence of motile diatoms and small cyanobacteria in the last added group that fill the porous space between larger and filamentous cells within the periphyton matrix. Another possible explanation for the increased density could be related to changes in the EPS composition during periphyton development. Indeed, different microbial species synthetize different EPS, which in turn determine the microenvironment of the cells within the biofilm by affecting porosity, density, water content, hydrophobicity, charge, sorption properties, cohesiveness, and mechanical stability [43]. For instance, it has been shown that biofilms dominated by taxa producing very low amounts of EPS are highly dense [44]. Overall, these our results provide additional detail on how physical structure parameters change during periphyton establishment and on the contribution of each phototrophic group and potentially their extracellular matrix to these changes. Further investigations are needed to address the contribution of EPS to the 3D structure of biofilms by analyzing their composition.

**Fig. 3.**
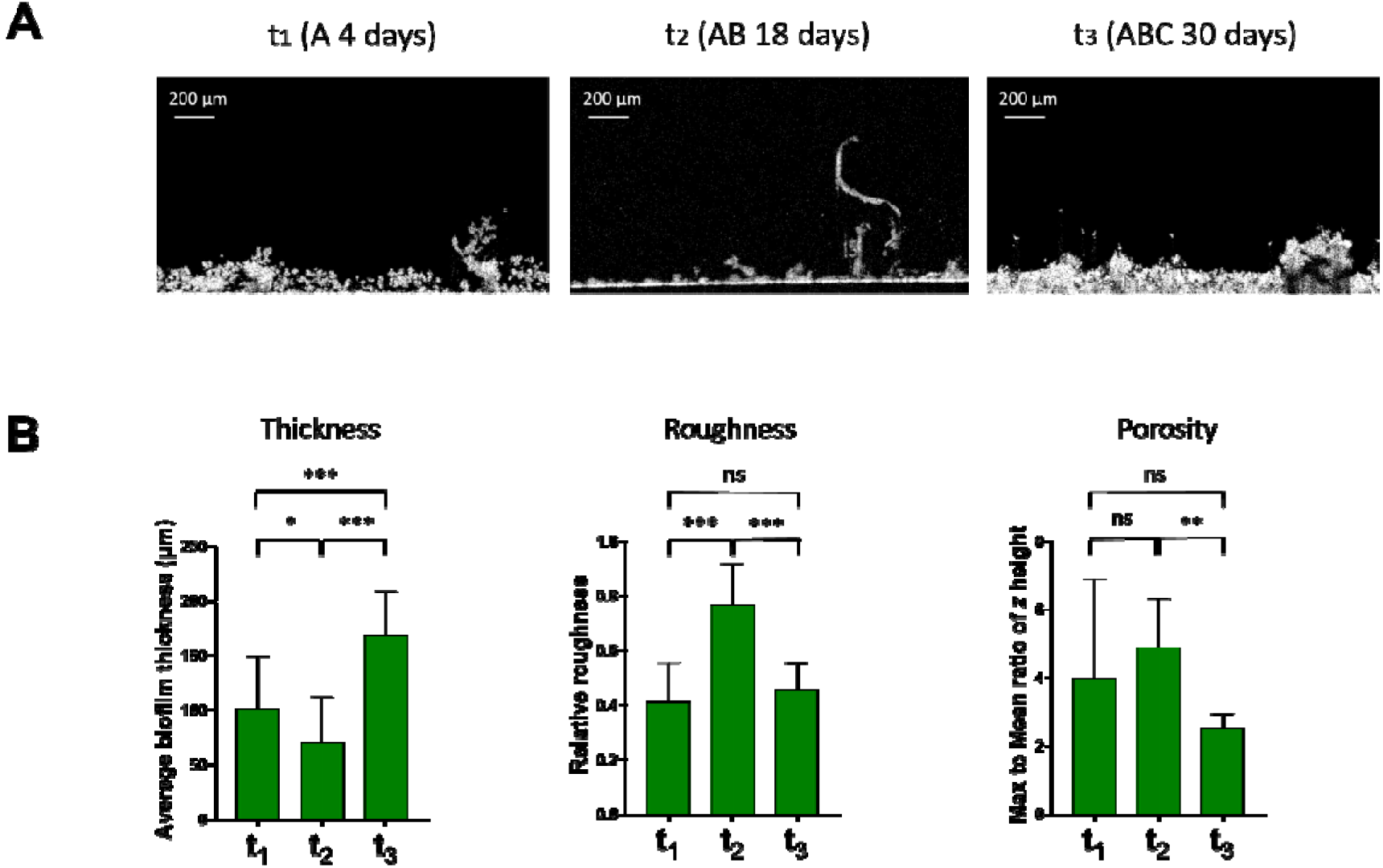
Changes of the periphyton physical structure during its establishment. Shown are representative OCT images obtained over 30 days of periphyton growth at time points t_1_ (A 4 days), t_2_ (AB 18 days) and t_3_ (ABC 30 days) **(A)**. White pixel: Biomass accumulation, black pixel: absent/low biomass accumulation. Scale bar: 200 μm. Mean thickness, surface topology (relative roughness) and internal porosity (mean to max ratio of *z*, where *z* is the height of the biofilm) were determined from 20-30 images and 5 biological replicates per time point **(B)**. Error bars represent standard deviations. * P<0.05, ** P<0.005 *** P<0.0001, post hoc Tukey test.

### Microbial diversity and composition during periphyton establishment

Analyses via confocal laser scanning microscopy (CLSM) of the intact periphyton at 30 days showed the presence of 12 to 15 different species that are co-existing in close proximity within the community (Supplementary Fig. 5). CLSM is a powerful technique to examine three-dimensional structure of the periphyton at the micrometer scale [45, 46]. However, it does not allow precise identification of species with similar cellular morphologies (e.g., small and round or long filaments) or provide quantitative information on the relative abundance of each genotype within the community. To overcome these limitations, we performed high-throughput amplicon sequencing of 18S rRNA (green algae and diatom), *rbcL* (higher taxonomic resolution for diatoms) and 16S rRNA (cyanobacteria) genes (see Methods and Supporting Information for more details). For this, we created our own reference database for the twenty-six single species that were used to establish the synthetic periphyton with their respective amplicon gene sequences (Supplementary Table 5). We also validated the accuracy of the amplicon sequencing approach for single species identification and quantification by using a mock community (see Supplementary Discussion, Supplementary Fig. 6 and Supplementary Table 6 for more details). Among the 26 species used to establish the periphyton, we were able to accurately identify 21 eukaryotic (diatoms and green algae) and 4 prokaryotic (cyanobacteria) species, with the only limitation to distinguish between *Pediastrum duplex* from *Pediastrum boryanum*.

Once the robustness of our approach was validated and single species were taxonomically assigned, we monitored community composition in the mature synthetic periphyton as well as changes over time during periphyton formation. In comparison to morphological analysis by CLSM, amplicon sequencing provided higher precision for single species identification. Indeed, characterization of the community composition revealed a diverse and complex mature periphyton composed of at least 22 different phototrophic species, corresponding to 13 diatoms, at least 5 green algae and 4 cyanobacteria, with various relative abundances (Fig. 4 A-C). For instance, *C. meneghiniana, A. pyrenaicum, S. acuminatus* and *S. vacuolatus* were the most abundant in the established periphyton at t_3_, representing approximately 75% of the final community (Fig. 4A). Moreover, all four cyanobacteria species that were introduced into the community were also detected in the mature periphyton (Fig. 4C and Supplementary Fig. 7D). The highest abundances were measured for *C. polonicus* and *M. glauca*, and the lowest one for *Phormidium sp*. These findings are in line with previous studies that identified these phototrophic species in either natural *in-situ* periphyton or periphyton grown in microcosms [11, 47, 48], demonstrating the ecological relevance of our synthetic microbial community.

**Fig. 4.**
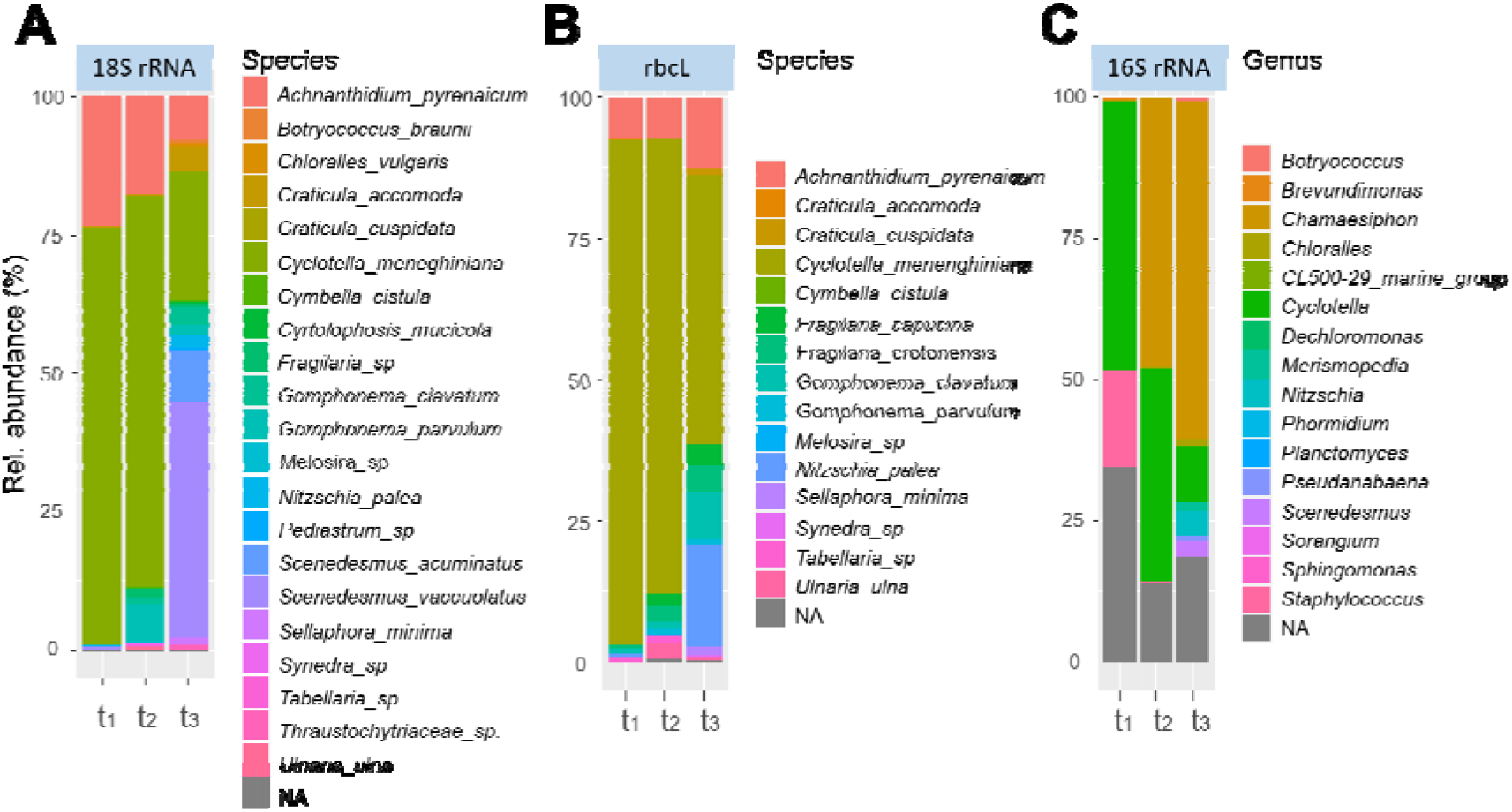
Genetic composition of the synthetic periphyton at the species level. Community composition profile and relative abundances (average; n = 9) were inferred from taxonomy-based clustering at species level of assigned 18S rRNA **(A)** and *rbcL* **(B)** gene sequences and at genus level of assigned 16S rRNA **(C)** at 4 (t_1_), 18 (t_2_) and 30 days (t_3_) during periphyton formation. NA: not assigned species.

Our results also show that the development of periphyton follows a succession of changes that are potentially governed by species interactions, as it has been previously reported [8, 9, 24, 49]. For instance while, as expected, only *C. meneghiniana* and *A. pyrenaicum* were detected at t_1_ (Fig. 4A), at 18 days (t_2_) after addition of the nine high profile diatom species from group B more species were detected (Supplementary Fig. 7A and 7E). The abundance of *C. meneghiniana*, on the other hand, decreased significantly at day 30 (t_3_) in the mature periphyton (Fig. 4A and 4B, Supplementary Fig. 7A and 7E), suggesting that the presence and/or the activity of group C species may have suppressed its growth. Note that the highest abundance of *C. meneghiniana* and *S. vacuolatus* in the established community at t_3_ might also be the result of the high gene copy number and/or PCR amplification efficiency with 18S rRNA primers for these species (see Supplementary Discussion and Supplementary Fig. 6A). Furthermore, *Tabellaria sp*. and *Melosira sp*. from group B, as well as *N. vermicularis* introduced with group C, were no longer detected in the mature periphyton, neither with 18S (Fig. 4A, Supplementary Fig. 7B and 7C, respectively) nor with *rbcL* (Fig. 4B, Supplementary Fig. 7F and 7G, respectively), probably as a result of species competition. Therefore, from twenty-six sequentially introduced phototrophic species, three species were no longer detected in the mature periphyton.

Once the workflow to establish the synthetic periphyton was set, we tested its reproducibility. For this purpose, we performed two independent experiments, with two months gap in between, each time starting with planktonic cultures of the single species and following the established workflow (summarized in Fig. 1). When comparing the experiments, our data on genetic diversity shows the reproducibility of the obtained community (Supplementary Fig. 8). Despite slight differences in relative abundances, especially at t_1_, community composition of the mature periphyton at day 30 was well conserved. Most importantly, species succession during periphyton establishment was also similar between both experiments and among biological replicates (n = 9). Taken together, these results demonstrate that by following a rational design and applying existing knowledge on microbial dynamics during periphyton formation we were able to obtain a reproducible and diverse synthetic community.

### Case study: single and combined effects of the herbicide terbuthylazine and increased temperature on periphyton structure and function

Understanding how the individual and combined effects of multiple environmental stressors impact species interactions is a key challenge. In complex microbial communities, such as periphyton, species have a large degree of freedom for interactions that further complicates the evaluation of the outcomes. By using the established synthetic periphyton as a model system, we aimed here to examine precisely how single species within the community and the community as a whole respond during periphyton development to the single and combined exposures to two major environmental stressors: the widely-used herbicide terbuthylazine at 0, 1, 10 and 100 nM and a change in temperature from 17°C to 20°C (see Methods for more details on the experimental design). Climate warming is expected to affect the composition of periphyton through either direct effect on the physical properties of the water column [50–52] and/or indirectly by affecting light availability and nutrient levels [52]. Our results on community composition during periphyton establishment provide strong evidence that increased temperature significantly impacted microbial diversity and altered species succession in the community (see Supplementary Table 12 for statistical analysis). When comparing between community composition profiles in the mature periphyton at 17°C and 20°C at the phylum level, the average abundances of green algae (Chlorophyta) and cyanobacteria decreased from 61% to 25% and from 65% to 49%, respectively, while the one for diatoms (Bacillariophyta) increased from 37% to 74% (Supplementary Fig. 9A and 9B). A detailed analyses at the single species level shows that the abundances of the green algae *C. vulgaris, S. acuminatus* and *S. vacuolatus*, which dominated the community at 17°C, strongly decreased with increased temperature (Fig. 5C and Supplementary Fig. 9C). Interestingly, the growth of these green algae as single species in planktonic mode is favored at 20°C than at 17°C [53, 54], suggesting the potential influence of complex species interactions when taxa grow within benthic communities. Among cyanobacteria, only the abundance of *C. polonicus* was negatively regulated by the elevated temperature (Fig. 5D and Supplementary Fig. 9E). In sharp contrast, several diatoms, such as *C. meneghiniana, C. cistulla, Tabellaria sp*. and *Melosira sp*., were significantly favored by the temperature increase (Fig. 5A and 5B and Supplementary Fig. 9C and 9D). These findings are in line with previous field studies, which reported on increased abundance of *Cyclotella, Melosira* and *Tabellaria* with increasing water temperature [51, 55]. Interestingly, *C. meneghiniana* and *Melosira* sp. were not impacted by the temperature increase at t_2_ (Fig. 5A and 5B, respectively). On the other hand, at t_3_ upon addition of group C species, their abundances significantly decreased at 17 °C but increased at 20 °C (Fig. 5A and 5B, Supplementary Fig. 9C and 9D, and Supplementary Table 12). Such results suggest that temperature-sensitive species from the group C potentially balance the growth of *C. meneghiniana* and *Melosira sp*. at 17 °C. However, when their own growth is reduced at 20 °C, they probably can no longer compete, allowing *C. meneghiniana* and *Melosira sp*. to grow within the community regardless of their specific tolerance to temperature. However, the contribution of heterotrophic bacteria, which might be favoured by increased temperature [56], cannot be excluded [57, 58]. Indeed, species-specific interactions between the algal host and associated bacteria might change under higher temperature and therefore impact microbial succession of phototrophs in periphyton. Additional characterization of the associated bacterial composition to each phototrophic species in the synthetic periphyton, as well as measurements of changes in the C:N:P molar ratio in periphyton for example [50], would provide a deeper insight into such interactions.

**Fig. 5.**
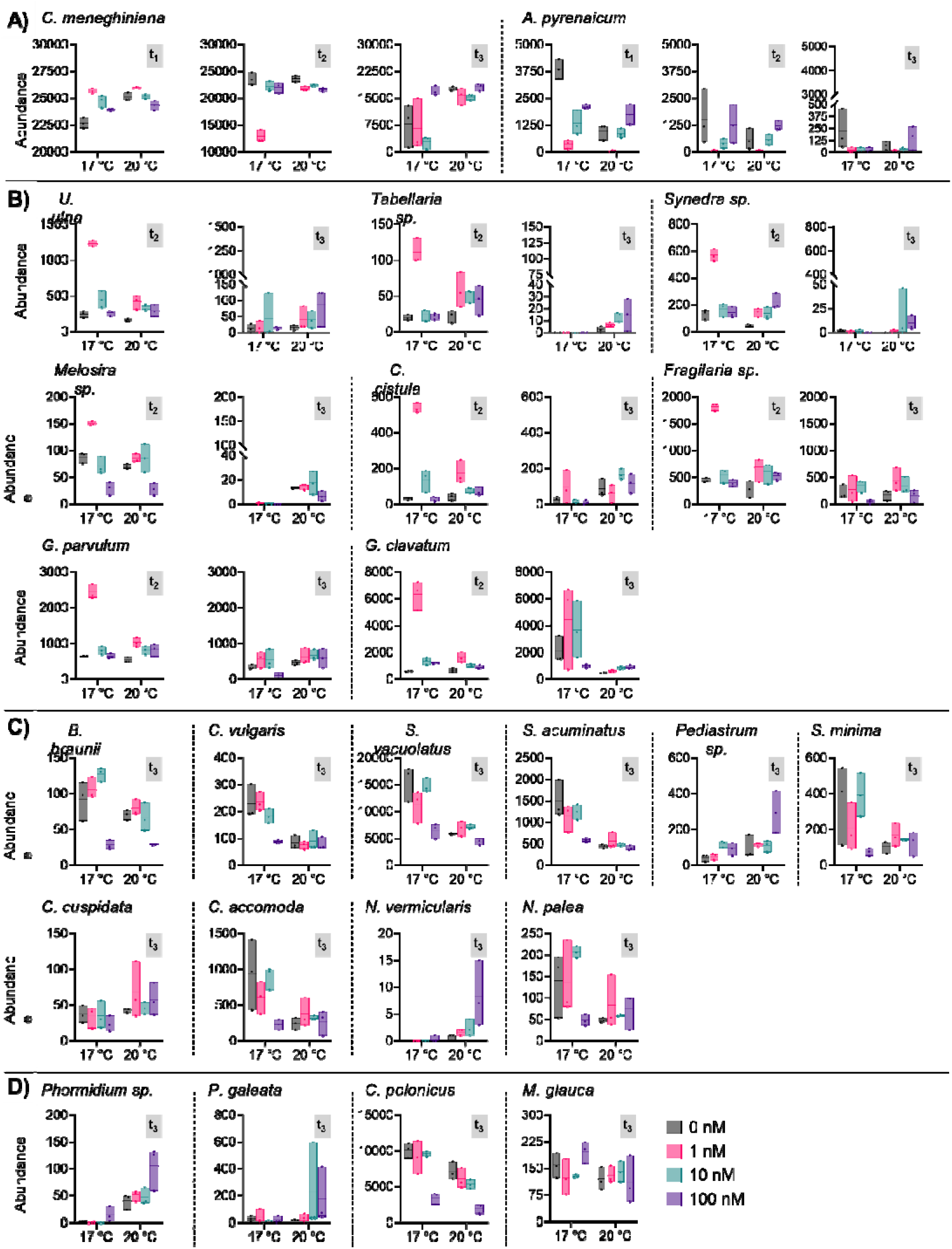
Abundance of single species during periphyton establishment. Shown is abundance of group A **(A)**, group B **(B)** and group C **(C)** eukaryotic species as well as group C prokaryotic species **(D)** measured at t_1_ (A 4 days), t_2_ (AB 18 days) and t_3_ (ABC 30 days) and grown under different conditions. The conditions correspond to 4 levels of the herbicide terbuthylazine (0, 1, 10 and 100 nM) and 2 different temperatures (17 and 20 °C). Horizontal lines in the black boxes correspond to the average values and the lower and higher limits of the standard deviation (n = 3). Results of the pairwise comparison (post hoc Tukey test) among all treatments for each species are shown in Supplementary Tables 13-15.

Exposure to increasing concentrations of terbuthylazine also led to changes in the microbial composition of the mature periphyton (Fig. 5 and Supplementary Fig. 9). The data on single species abundances shows that the large majority of green algae and some diatom species were negatively affected only by the highest terbuthylazine concentration (Fig. 5 and Supplementary Fig. 9). On the contrary, the relative abundance of the low profile diatom *C. meneghiniana* (Fig. 5A), green algae *Pediastrum sp*. (Fig. 5C) and the cyanobacterium *Phormidium sp*. (Fig. 5D) increased significantly (Supplementary Table 12) upon exposure to 100 nM terbuthylazine. This reflects their competitive advantage under unfavorable conditions for most of the other species within the community. Such findings are in line with those of previous studies that reported the presence of these species in natural periphyton that had been exposed to micropollutants such as herbicides [11, 59, 60], indicating their ability to tolerate high levels of pollutants. It has been previously reported that toxicity of PSII inhibitor herbicides does not depend on the 3D structure or biomass of biofilms but rather on the physicochemical characteristics of the substance itself, such as hydrophobicity (water solubility) and lipophilicity (fat solubility) [61–63]. The tolerance differences towards terbuthylazine we found among the species may be therefore related to variable capability of microbial species within the community to implement defense mechanisms towards herbicides, including physiological acclimatization related to the uptake, translocation, degradation and excretion of the herbicide [64]. The EPS composition, which contains a significant amount of proteins, can also provide additional sites for interactions with the nonpolar or lipophilic regions of the herbicide and thus interfering with its bioavailability and toxicity [65, 66]. Further experiments are needed to specifically investigate such toxicity and adaptive cellular mechanisms in each single species, as well as the influence of EPS composition.

As conceptualized by the subsidy-stress concept, environmental factors, such as increased temperature, can be beneficial for microorganisms and consequently mitigate the negative effects of micropollutants [67]. This is in line with our results on the relative abundances of single species, which show significant beneficial interaction between temperature and terbuthylazine (Supplementary Table 12). Indeed, the negative effects of the highest concentration of terbuthylazine in the mature periphyton at 17°C were no longer observed at 20°C for several diatoms (e.g., *A. pyrenaicum* at t_1_, *Synedra sp*. at t_3_ and *G. parvulum* at t_3_,) and green algae (e.g., *S. vacuolatus*, *S. acuminatus* and *C. vulagaris* at t_3_) (Fig. 5). Interestingly, some of the high profile diatom species such as *Synedra sp., Tabellaria sp*. and *C. cistula*, were not at all affected when exposed to terbuthylazine or temperature alone (Supplementary Table 12). Clearly, such results confirm that multiple interactions among species and temperature occur, circumstances that complicate the assessment of risks caused by chemicals in complex field situations. Because species have different temperature optima for growth but also variable sensitivities to chemicals, such interactions can now be assessed and better understood under controlled conditions with our synthetic periphyton model. This can be achieved by examining responses of the single species and in the community to temperature and herbicide gradients.

One additional insight from our case study is that both increased temperature and exposure to the herbicide terbuthylazine affected negatively the 3D-structure and biomass but not the PSII quantum yield of the periphyton. Despite the observed growth during periphyton development in all treatments, the overall periphyton 3D-structure (Supplementary Fig. 10), thickness (Supplementary Fig. 11A) and concentration of attached cells to the colonization substrate (Supplementary Fig. 12A) were significantly lower at 20°C compared to 17°C. Such results can be explained by the sharp decrease in the relative abundances of most of the green algae and diatom species composing group C upon increased temperature or exposure to terbuthylazine (Fig. 5, Supplementary Fig. 9). This conclusion is also supported by the fact that these negative effects were mainly observed after addition of group C species, confirming their important contribution to the benthic biomass and structural development of the periphyton. Despite the clear effects on benthic biomass and physical structure, but also microbial diversity and composition, exposure to terbuthylazine and increased temperature did not lead to notable effects at the functional level (Supplementary Fig. 13). PSII quantum yield was similar in all treatments and time points, except for periphyton exposed to 100 nM terbuthylazine at 18 days (i.e., before addition of group C). Overall, these results from our synthetic periphyton are in line with previous observations [19, 68] and might be related functional redundancy with regard to photosynthetic function when less sensitive species of the same functional group compensate for the loss of other species in order to ensure important community functions [69]. However, further specifically designed studies are needed to clearly demonstrate functional redundancy.

## Conclusion: potential applications and limitations

In order to answer important ecological questions, such as how species interactions can translate into complex community-level processes and properties, we developed a workflow for a reproducible establishment of a synthetic freshwater periphyton. While the majority of previously established synthetic microbial communities consist of only two to four genotypes or focused only on few diatom species, our synthetic periphyton consists of at least 22 different microbial phototrophic species, growing and interacting in close proximity. Because of its reproducibility, the synthetic periphyton that we established offers great opportunities for theory testing through targeted manipulation and for exploring crucial ecological processes, such as diversity-function relationships or resistance and resilience to environmental stressors. This can be achieved via the elaboration of study designs in which the inoculum composition (i.e., species pool), species densities, colonization durations and order or manner of introduction, are manipulated. For instance, all taxa can be introduced at once into the growth chambers, instead of sequentially, and microbial succession can be monitored at the single-species level. This will allow testing the influence of mass effects from dispersal versus competitive interactions associated to a given environment on succession, by comparing the outcomes with those of sequential addition. By knowing the exact composition of the synthetic community, studies comparing between the responses of microbial species growing alone or when associated in a biofilm, where multiple interactions occur, can be performed with any abiotic or biotic stressor. Species interactions, in the presence of stressors or not, could be also systematically examined through a combinatorial approach in which one or several taxa are excluded from the species pool. Finally, although our focus was on phototrophs, future studies could also consider introducing other groups of organisms, e.g. heterotrophic bacteria or (micro)grazers, in order to examine, for example, trophic networking.

Despite the numerous advantages and potential applications for the established synthetic community, it also has some limitations that need to be considered for future investigations. In this study, the synthetic communities were established under specific environmental condition (e.g. high nutrient concentration and specific actinic light exposure). Under another environmental filtering, community development and species succession might be therefore different [8, 9, 70–72]. The absence of flow, as is the case in our experimental system, might also lead to a potential formation of gradients of light, oxygen, pH and nutrients within the biofilm [72, 73], which in turn can affect the biofilm development, extracellular matrix production, species interactions and competition [74, 75]. Developing the synthetic periphyton in sterile flow-through chambers [76], would mimic to some extend conditions in streams and rivers. Such system will also offer more flexibility in terms of growth medium volume and will allow testing effects of different flow levels. Finally, despite their importance in natural periphyton, filamentous green algae (e.g. Ulothrix, Mougeotia and Oedogonium) were excluded from the pool of species due to the difficulties in quantifying their exact cell number in the initial inoculum. The latter being a crucial criterion for reaching a high level of reproducibility.

## Methods

### 2.1. Cultures of the phototrophic species

Non-axenic planktonic cultures of the 34 selected single phototrophic species were obtained from (1) SAG Culture collection of algae (Goettingen university, Germany), (2) CCAC Culture collection (University of Cologne, Germany) and (3) TCC Thonon culture collection (INRAE, France) (Supplementary Table 1). Because several taxa require different optimal temperature and light intensity for their growth, all species were adapted for six months to grow in the same freshwater medium COMBO (see Supplementary Table 2 for the detailed composition) [77] at 17°C and photon irradiance of 40 to 60 μmol s^−1^ (Philips Master LED tube Value 1200 mm InstantFit 865, 16.5 W, MLT Moderne Licht-Technik AG, Wettingen, Switzerland), as measured with PAR Quantum sensor Spectro Sense 2 (Skye Instruments Ltd, UK), with a photoperiod of 16h light: 8h dark. For this, we progressively increased the light intensity and decreased the temperature for diatoms and green algae, respectively, until reaching the required conditions for our experimental design. Twenty-six out of the 34 species were successfully acclimatized (i.e., comparable growth to their optimal initial temperature and light condition) and used for the establishment of the synthetic periphyton. All main planktonic cultures were kept in triplicate in sterile 300-mL Erlenmeyer flasks containing 150 mL of COMBO and placed on a shaker operated at 30 rpm. Every 3 weeks, 50 mL of these cultures were centrifuged at 2000 g, the supernatant removed and cells in the pellet resuspended in 1 mL fresh COMBO. 200 to 500 μl of this suspension were added to 75 mL fresh COMBO for growth measurements and 500 to 1000 μl were added to 150 mL of fresh COMBO and kept as backup cultures.

Depending on the species morphology, growth rates and maximum reached biomass of each single species in suspension were generated from optical density and/or cell count measurements of the planktonic cultures over time. The flasks were incubated in the same conditions as described above and 2 mL were taken every 48 h for the growth measurements. Optical density of the suspensions at 685 nm was measured with a spectrophotometer (UVIKON 930, Kontron Instruments, Basel, Switzerland). Cell concentration was determined by using an automated CASY cell counter Model TT (Roche Innovatis AG) with 60 μm capillary, 2.18 μm - 30 μm evaluation cursor and 1.5 – 29.83 μm normalization cursor to take into consideration the different cell sizes. The obtained data were fitted in a logistic model as function of time in order to derive the growth rate constant (*k*) and maximum reached biomass for each species in suspension (detailed results are provided in the Supplementary Tables 3 and 4).

### 2.2. Experimental system and design for the establishment of the synthetic periphyton

The establishment of periphyton in our study was performed under static conditions in sterile 1-well Nunc™ Lab-Tek™ II Chamber Slides (74 × 25 mm, Surface = 1850 mm^2^, Volume = 4 mL, Thermo Scientific), which consists of a removable polystyrene media chamber attached to a standard glass slide. First, the bacteria *Sphingomonas elodea* were inoculated from the glycerol stock into a sterile Erlenmeyer flasks containing 100 mL ATCC rich broth medium (8.0 g Nutrient Broth (BD cat 234000) in 1 L deionized water). After 48 h of growth at 200 rpm and 30 °C, optical density of the bacterial culture was measured at 600 nm. Chambers were then inoculated with 4 mL *Sphingomonas elodea* in ATCC rich broth medium with an OD_600_ of 0.4 and incubated at room temperature under static conditions for bacteria to colonize the surface and produce the EPS matrix. Three days after inoculation, the medium was removed and chambers gently washed once with COMBO before adding a mixture of the two group A species at a one to one ratio in 4 mL COMBO. After 4 days of incubation, medium with non-attached cells was removed carefully and 4 mL of COMBO containing a mixture of the nine species that compose group B were introduced to the chambers. After another 14 days, the medium with non-attached cells was again removed before addition of the 18 species of group C in 4 mL COMBO (see Table 1 and Supplementary Table 2 for details). Finally, the chambers were incubated for another 12 days, leading to a total duration of 30 days for periphyton establishment, starting from the introduction of group A. In order to avoid potential effects of nutrient limitation, 2 mL of COMBO were carefully replaced in the chambers every second day. All incubations were performed in the same temperature- and light-regulated room as the planktonic cultures (i.e. 17°C and photon irradiation of 40 to 60 μmol s^−1^, photoperiod of 16h light: 8h dark). Details about the experimental design, total number of inoculated chambers and specific measurements are provided in Supplementary Information (Supplementary methods, Supp. Fig.1).

### 2.3. Periphyton characterization

#### 2.3.1. Surface colonization area

Surface coverage of the different phototrophic species during periphyton establishment was checked microscopically at low magnification. At each sampling time point the supernatant (4 ml) was removed, chambers were washed with 4 ml COMBO medium and surface coverage of surface-attached cells suspended in 4 mL fresh COMBO media was measured (Supplementary Fig. 1). Imaging was performed with an inverted Leica microscope (Plan Neo Fluar Z 1.0□×/0.25, FWD 56□mm objective) equipped with a CCD camera (Axiocam 506 mono, Zeiss). Fiji an open-source image processing software based on ImageJ (version 1.53c) was used for image analysis [78]. Note that since biofilm observations were done without z-stacking, the surface coverage corresponds to the biofilm coverage of the chamber bottom. Twenty-seven 9□×□3 tile acquisitions were acquired, montaged and stitched together, covering large area of the chamber surface (10□× magnification, 1.2□μm□pixel^−1^ resolution). A montage of images and analysis were performed by using custom-written macro in ImageJ (version 1.53c). In brief, after thresholding based on automated algorithm “Huang dark” [79], a counting mask was generated and surface-attached particles were analyzed and counted. The script for the ImageJ using custom-written macro and “Huang dark” auto-thresholding is provided in the Supplementary Methods. An average value from microscopy of three to five chambers was calculated per time point and condition.

#### 2.3.2. Cell density and cell number

To quantify density or number of cells within the periphyton (attached to the surface) and in suspension (supernatant) during periphyton establishment, total supernatant (4 mL) was collected and the attached cells were suspended in 4 mL fresh COMBO media and scratched from the entire glass surface of the chambers. Cell suspensions of attached cells were diluted 1:20 and 1:400 for OD_685_ and CASY cell count measurements, respectively. Three to five chambers (biological replicates) were retrieved per time point and condition. A minimum of three measurements were performed per replicate.

#### 2.3.3. Effective photosystem II quantum yield measurement

After removing the supernatant from the chambers and adding 4 mL fresh media, the effective photosystem II quantum yield, (Φ_PSII_) of all biofilms was assessed by chlorophyll *a* fluorescence measurement, as a proxy for photosynthetic activity [11, 80, 81]. Measurements were performed with an Imaging Pulse Amplitude Modulated fluorimeter (MAXI-I-PAM, M-series, Heinz Walz GmbH, Germany) equipped with an IMAG-K CCD camera and red (660 nm) and near-infrared (780 nm) LEDs. The average of the whole field of view (i.e. the entire chamber slide area of 1850 mm^2^) was used to obtain the effective PSII quantum yield, which was automatically calculated by the machine software ImagingWin. I-PAM measurements were always performed at the same daytime for all experiments (i.e. 4 hours after the start of the light cycle in the incubation room) and at room temperature. Every biofilm was measured with a Photosynthetic Active Radiation of 81-μmol□m^−2^□s^−1^ with an actinic light (LED array of the I-PAM), which was about the same light irradiance as the levels used for the growth of the single species and biofilms. The duration of actinic light exposure in the I-PAM was optimized before the start of each measurement, with constant yields being observed after adaptation times exceeding 2.5□min. We also qualitatively checked that applied actinic light was in the detection range; above the light limiting and below the light saturating conditions, by derived rETR versus irradiance curves. Chlorophyll-a from each periphyton was excited at 665 nm and measured at 780 nm after applying a single light saturation pulse (2800 μmol□m^−2^□s^−1^) to calculate the effective PSII quantum yield (□’) as:

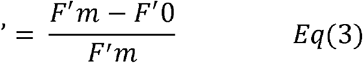

where *F*′*_m_* is the maximum fluorescence after the saturation pulse and *F*’0 is the steady-state fluorescence.

#### 2.3.4. 3D-structure of periphyton

##### 2.3.4.1. Optical Coherence Tomography (OCT)

Optical Coherence Tomography (Ganymede model 930 nm Spectral Domain, Thorlabs GmbH, Dachau, Germany) with a central light source wavelength of 930 nm was used to investigate the meso-scale structure of periphyton. The use of long wavelength light allows to penetrate up to a depth of 1.9 mm with axial and lateral resolutions of 4.4 μm and 15 μm, respectively. 20-30 scans (XZ pane image) of 2×1 mm were acquired per replicate at t_1_ (at day 4, just before addition of group B species), t_2_ (at day 18, just before addition of group C species) and t_3_ (at day 30, the end of periphyton establishment) during periphyton development. Three replicate samples per timepoint (and condition) were examined. Image analysis software developed under Matlab® (MathWorks, Natick, US) was used to analyze OCT image. The Matlab script is available on request. Image analysis consisted of the following steps: (1) surface-biofilm interface detection (grey-scale gradient analysis); (2) image binarization (automatic threshold selection); (3) calculating physical properties of the biofilm. Biofilm thickness was calculated based on the number of pixels found between the top edge of the periphyton and the upper chamber surface of each OCT image, using a pixel scaling factor to obtain the biofilm thickness in microns.

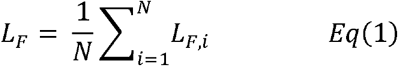

where L_F,i_ (μm) is the biofilm thickness from a single A-scan (i.e., light reflected from each optical interface) in the corresponding B-scan (i.e., cross-sectional reconstruction of a plane from a series of A-scans across the structure) and N is the total number of A-scans.

The relative roughness coefficient (-) of the biofilm was calculated according to [82].

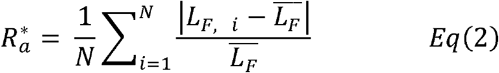

where *i* represents an A-scan and the overall number of A-scans. Biofilms with a smooth surface and only a few variations from the mean biofilm thickness have low values close to 0. The higher the roughness coefficient, the more variations are expected from the biofilm surface [83]. The roughness coefficient is based on the individual thickness measurement L_F,i_, the sample mean thickness L_F_, and the number of thickness measurements N.

OCT images does provide some indication of how dense or loose a biofilm is structured, and fraction of light/dark pixels or greyscale distribution can be quantified. Wagner *et al*. [84] suggest zones of low signal intensity represent areas of absent biomass (i.e., voids, pores). The distinction between solid materials and voids relies on the assumption that absence of OCT signal equals voids. However, such an assumption and the resulting porosity quantification is debatable.

##### 2.3.4.2. Confocal laser scanning microscopy

At day 30, confocal laser scanning microscopy (CLSM) was used to visualize the periphyton structure at the μm-resolution and to detect single species within a dense 3D-structure. For this purpose, polystyrene media chambers of 1-well Nunc™ Lab-Tek™ II Chambers Slides were removed with special adaptors and glass slides placed into a petri dish with 25 mL fresh COMBO medium (see Supplementary Information for more details). Samples were examined with upright Leica SP5 confocal laser scanning microscopy (Heidelberg GmbH, Mannheim, Germany). Algae and cyanobacteria were detected through the autofluorescence of chlorophyll-a signal (excitation 630 nm, emission 650-720 nm). All observations were performed using a 40×/0.75 NA objective (Leica, Plan-Apochromat®). Periphytons were scanned at 2-μm intervals in z, starting at the glass surface, resulting in a series of individual images at different depths (*z*-scan) and an XY size of 1024×1024 with zoom factor of one. The typical z-range covered 200-300 μm. CLSM data was analysed by Imaris 9.3.0. Three replicate samples were examined per condition. For image analysis, three 3D-images per replicate were recorded, producing nine 3D-images per periphyton sample.

#### 2.3.5 Next generation sequencing for microbial community composition

##### 2.3.5.1 DNA extraction, library preparation and sequencing

In order to compare the composition of prokaryotes (i.e. cyanobacteria) and eukaryotes (i.e. diatoms and green algae) in periphyton, total genomic DNA was extracted from an aliquot of 1 mL from each periphyton suspension at t_1_ (4 days), t_2_ (18 days) and t_3_ (30 days) during the experiment. Five biological replicates from the experiment 1 and four biological replicates from the experiment 2 (total n=9) were used to obtain taxonomic composition of the community at the indicated time point. The samples were centrifuged at 14’000 *g* for 30 min at 4 °C and the pellets stored at −20 °C until their analyses. DNA extraction was performed by using the DNeasy Plant Mini Kit (Qiagen) following the manufacturer’s instructions. Prior to DNA extraction, physical disruption of cells was performed using cooling BeadRuptor Elite (OMNI International, US) and bead tubes from the DNeasy PowerBiofilm Kit (0.5 mm glass, 0.1 mm glass and 2.8 mm ceramic beads). Total DNA was then quantified with a Qubit (1.0) fluorimeter following the recommended protocol for the dsDNA HS Assay (Life Technologies, Carlsbad, CA, USA).

Library construction consisted in a two-step PCR process (see Supplementary Methods for more details). The first PCR amplified the 16S rRNA from cyanobacteria, V4-V5 region of the 18S rRNA gene for eukaryotes and *rbcL* gene using two different primer sets with overhang adapters from [85], [86] and [87], respectively (Supplementary Table 8). The first PCR was performed in triplicate for each DNA sample, negative PCR controls, as well as positive PCR controls for 16S rRNA, 18S rRNA and *rbcL*, consisting of mock communities (Supplementary Tables 5, 9-11). The second PCR, consisting in a limited-cycle amplification, was carried out to add multiplexing indices and Illumina sequencing adapters. The libraries were then normalized and pooled to achieve a 3.2 nM concentration. Paired end (2 × 300 nt) sequencing was performed on an Illumina MiSeq (MiSeq Reagent kit v3, 300 cycles) following the manufacture’s run protocols (Illumina, Inc.). The MiSeq Control Software Version 2.2, including MiSeq Reporter 2.2, was used for the primary analysis and the demultiplexing of the raw reads. All raw sequences are available at the National Center for Biotechnology Information (NCBI) under the SRA accession ID PRJNA764864.

##### 2.3.5.2 Sequencing data processing, amplicon sequence variants binning and taxonomic assignment

The reads were checked for quality and end-trimmed by using FastQC v0.11.2 [88] and seqtk (https://github.com/lh3/seqtk), respectively. For 16S rRNA and *rbcL*, the reads were merged using FLASH v1.2.11 (minimum and maximum overlap of 15 and 300 bp, respectively; maximum mismatch density of 0.25) [89] while only reads obtained with the forward primer were considered for 18S rRNA. The primers were trimmed by using cutadapt v1.12 (wildcards allowed; full-length overlap; error rate 0.01) [90]. Quality filtering was performed with PRINSEQ-lite v0.20.4 (minimum quality mean 20; no ambiguous nucleotides; dust low-complexity filter with a threshold of 30) with a subsequent size and GC selection step (size selection range 330–600 bp; GC selection range 30–70%) [91]. The reads were processed with an Amplicon Sequencing Variants (ASV) analysis [92]. The sample reads were first denoised into ASVs with UNOISE3 in the USEARCH software v. 11.0.667. The final predicted taxonomic ASV assignments were performed with Mega6 based on our own sequence database (see 2.3.5.3. and Supplementary Discussion) by applying a maximum likelihood phylogenetic tree using nearest-neighbor interchange (Supplementary Table 6). The total reads obtained at each step of bioinformatic filtration are reported in Supplementary Tables 9-11.

##### 2.3.5.3. Generation of single species DNA barcodes database

In order to accurately assign species in a sample we created our own DNA barcode reference database containing all twenty-six single species that were used to establish the synthetic periphyton and their respective barcodes (Supplementary Table 6). For this purpose DNA from single species grown in planktonic mode was extracted as described in 2.3.5.1. The corresponding genes (18S rRNA, *rbcL*, 16S rRNA genes) were PCR amplified form the extracted DNA using primers and conditions listed in Supplementary Table 7. PCR purified fragments were sent for Sanger sequencing. Sequences were quality trimmed using SnapGene 5.3.2 and initially aligned using ClustalW as implemented in Mega 6. Sequences were deposited in GenBank under accession numbers OL304135-OL304156 (18S rRNA), OK669058-OK339073 (*rbcL*) and OK737797-OK737800 (16S rRNA).

### 2.4 Case study: assessing the single and combined effects of the herbicide terbuthylazine and increased temperature on the synthetic periphyton structure and function

Terbuthylazine is a widely used photosystem II inhibitor herbicide, with environmental concentrations that can vary from 2 to 600 ng/L (corresponding to 0.08 - 2.74 nM) in Swiss streams that are impacted by wastewater effluents or agriculture [37, 93]. In the context of global warming, increasing temperature is considered as one of the most critical threats to biodiversity preservation and ecosystem functioning [94]. What is more, temperature can interact with micropollutants such as herbicides, which complicates the prediction of potential outcomes. Terbuthylazine has been shown to be toxic to freshwater algae, with EC_20_ and EC_50_ values ranging from 10 to 200 nM, depending on the species and measured endpoint [95–98]. In this context and as a proof of principle, we experimentally tested the single and combined effects of these two stressors based on a full factorial replicated (n = 3) design. Synthetic periphyton communities were established, as described previously, in the presence of four levels of terbuthylazine (i.e., 0, 1, 10 and 100 nM) and under two temperatures (17°C and 20°C). During the 30 days of periphyton establishment, 2 mL COMBO, supplemented or not with terbuthylazine, were exchanged every two days, to avoid nutrient limitation and maintain a constant exposure to the herbicide. Periphyton was sampled at days 4, 18 and 30 to measure benthic biomass, 3D-structure, photosynthetic yield and species composition, as described above.

### 2.5 Data analysis

GraphPad Prism 9 was used for statistical tests and visualization of data. Significant differences in phototrophic biomass, quantum yield, OCT and taxonomic abundance were assessed by one-way ANOVA during the periphyton establishment and two-way ANOVA in the case study to test for the interactions of the herbicide and temperature. When effects were significant, the ANOVAs were followed by Tukey’s post hoc tests. Normality and homogeneity of variance were checked prior to the ANOVA analysis (Kolmogorov-Smirnov’s and Levene’s tests, respectively). Data that were not normally distributed were log transformed. The significance level for all tests was set at α =□0.05.

Sequencing data analyses were performed with the R package Phyloseq version 1.32.0 (McMurdie and Holmes, 2013). After rarefaction, 16S, 18S rRNA and *rbcL* datasets were composed of samples containing the same number of reads (17209, 26097 and 2652 reads, respectively). Graphical representations were generated with the R package “ggplot_2_” or GraphPad Prism9.

## Supporting information

supplementary Figures, Methods and Discussion

## Data availability statement

All data generated or analyzed during this study are included in this published article (and its supplementary information files). MiSeq data that support the findings of this study have been deposited in the National Center for Biotechnology Information (NCBI) under the SRA accession ID PRJNA764864. Sanger sequences of single species were deposited in GenBank under accession numbers OL304135-OL304156 (18S rRNA), OK669058-OK339073 (rbcL) and OK737797-OK737800 (16S rRNA).

## Code availability statement

The custom-written macro for the ImageJ software, using the “Huang dark” autothresholding, is provided in the Supplementary Information. The Matlab script used to analyze OCT image with software developed under Matlab® (MathWorks, Natick, US) will be available upon request.

## Acknowledgements

This work was supported by the project ToxAdapt funded by Eawag. We thank Alexa Agatiello for the technical assistance in the DNA samples preparation for sequencing, as well as Dr. Louis Carles and Prof. Kristin Schirmer for the fruitful discussions and their feedback on previous drafts of the manuscript. We would also like to thank Silvia Kobel and Jean-Claude Walser (Genetic Diversity Centre (GDC), ETH Zürich) for their support with community composition analysis. Data on molecular diversity produced and analyzed in this paper were generated in collaboration with the GDC, ETH Zürich.

## Contributions

OL and AT conceived and designed the study. OL and BW performed the experiments. OL and ND performed the quantitative OCT analysis. OL and AT analyzed the data and wrote the first draft of the manuscript. All co-authors contributed to the data interpretation and subsequent revisions of the manuscript, and they approved the final submitted version.

