## supplementary Figures, Methods and Discussion for "Synthetic periphyton as a model system to understand species dynamics in complex microbial freshwater communities"

Department of Environmental Toxicology (Utox), Swiss Federal Institute of Aquatic Science and Technology (Eawag), Dübendorf, Switzerland.

Überlandstrasse 133

8600 Dübendorf

Switzerland

**Supplementary Methods**

*Experimental design and measurements*

The total number of chambers were inoculated at the start of each experiment and then used at for the specific measurements. At sampling time point the same set of replicates was used for all the measurements, except for the confocal laser scanning microscopy. Thus, for the establishment of the synthetic periphyton at least 20 chambers ((5 replicates x 3 time points) + (5 replicates for CLSM)) and for the case study at least 120 chambers ((8 conditions x 4 replicates x 3 time points) + (8 conditions x 3 replicates for CLSM)) were inoculated. All measurements except the confocal microscopy were performed at three sampling times: at t_1_ (4 days), t_2_ (18 days) and t_3_ (30 days). First, undisruptive measurements were performed (Supplementary Fig. 1A), followed by the disruptive measurements (Supplementary Fig. 1B), where supernatant and surface-attached cell fractions were analyzed:

(A) physical structure measurements using OCT and photosystem II quantum yield with iPAM were performed on intact periphyton;
(B) benthic biomass measurements with optical density and cell counter CASY, surface colonization area and microbial composition analysis with next-generation sequencing were performed on surface-attached biofilm after removal of the supernatant.

Confocal microscopy was performed at the end of periphyton establishment, at day 30 (Supplementary Fig. 1C). For this, three to five chambers per condition were taken, plastic walls of the chambers were removed with special adaptors and the glass slides were placed in petri dish with 25 mL fresh COMBO medium. Schematic representation of sample preparation for CLSM imaging is shown in Supplementary Fig. 1C. The typical z-range covered 200-300 μm. Live imaging was performed on intact biofilms without fixation or staining.

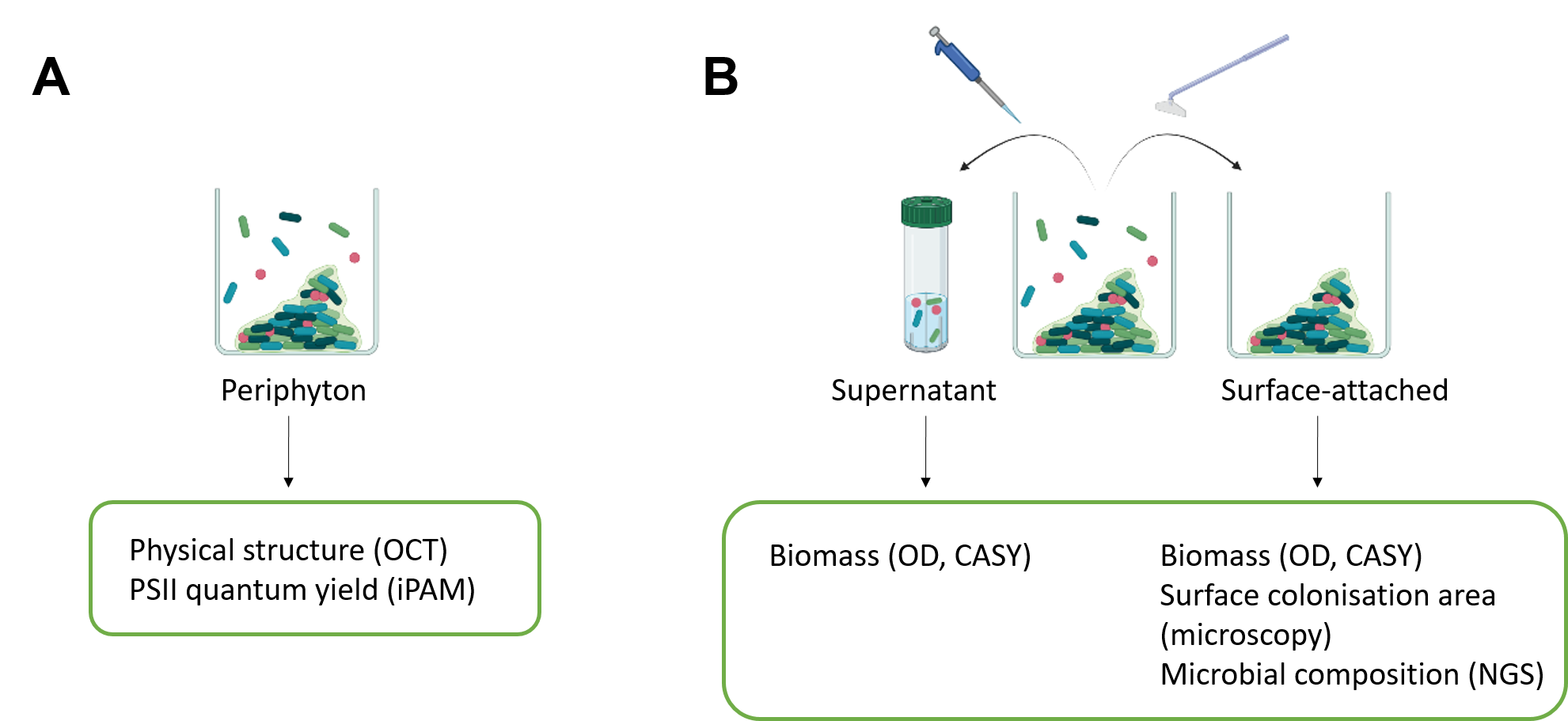

**Supplementary Fig. 1:** Schematic representation of the measurements performed during the establishment of the synthetic periphyton. Physical structure and PSII quantum yield measurements were non-destructive measurements **(A)**. Benthic (surface-attached cells) biomass, surface colonization area and microbial composition of the surface-attached cells were determined after the supernatant was removed **(B)**. Non-benthic (supernatant cells) biomass was also determined **(B)**. Confocal laser scanning microscopy (CLSM) imaging was performed at day 30 (t_3_) with the synthetic periphyton (N = 3 replicates) grown in 1-well Nunc^TM^ Lab-Tek^TM^ II Chambers Slides (**C-i**). For this purpose, by using specific adaptors **(C-ii)** the plastic walls of the chambers were removed **(C-iii)** and the glass slide was placed into the sterile petri dish with 25 ml COMBO medium **(C-iv)**. Microscopic observations were performed on at least five different points of one glass slide using upright Leica SP5 confocal laser scanning microscopy (Heidelberg GmbH, Mannheim, Germany) and 40×/0.75 NA objective (Leica, Plan-Apochromat®).

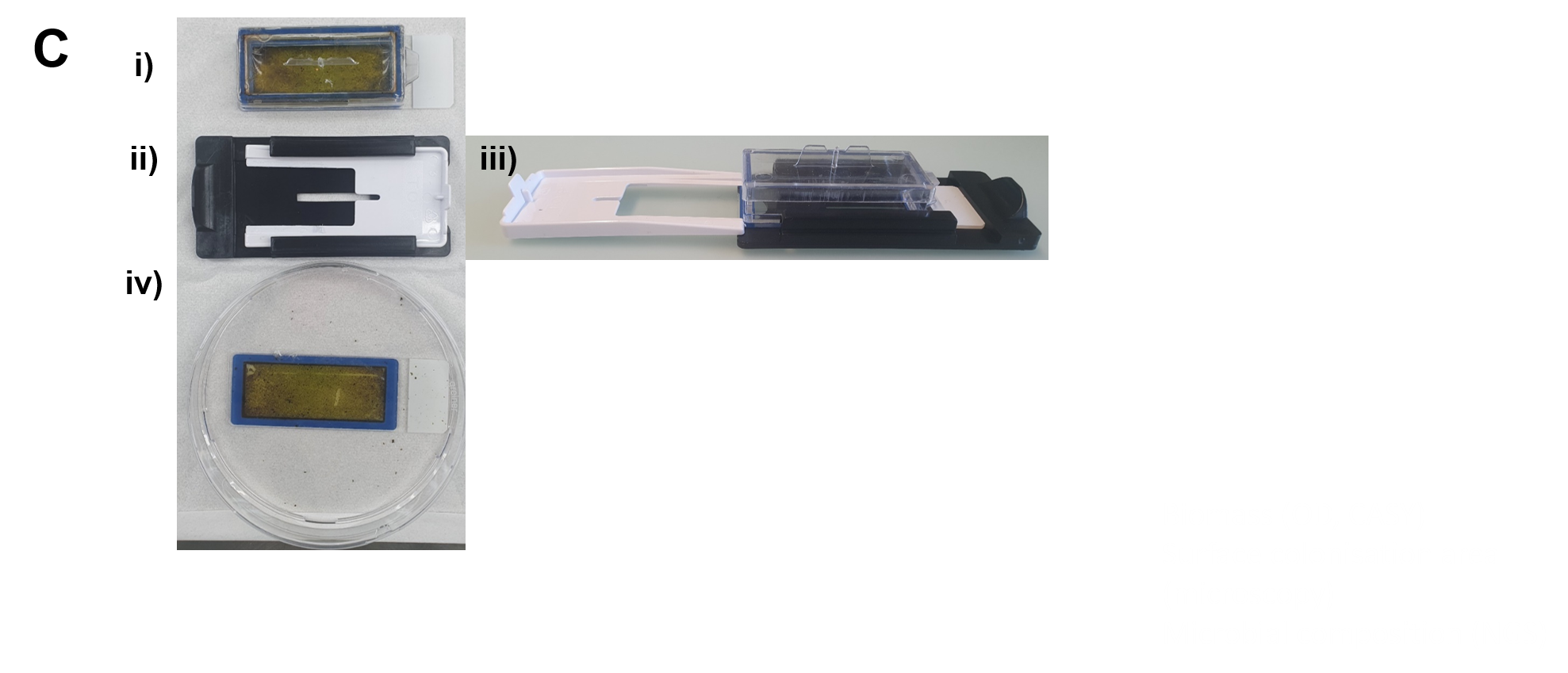

#### DNA Extraction, library construction and sequencing

Library construction consisted in a two-step PCR process. The first PCR amplified the 16S rRNA from cyanobacteria, V4-V5 region of the 18S rRNA gene for eukaryotes and *rbcL* gene using two different primer sets with overhang adapters from [1, 2] and [3] respectively (Supplementary Table 8). The first PCR was performed in triplicate for each DNA sample, negative PCR controls, as well as positive PCR controls for 18S rRNA, *rbcL* and 16S rRNA, consisting of mock communities (Supplementary Tables 9-11). The initial conditions of the first PCR, including cycle number, were first determined via quantitative RT-PCR following an internal protocol developed by the Genetic Diversity Center (GDC), Zürich. A minimal cycle number was used to get enough amplicons while limiting PCR-linked bias. Bias was further limited by performing the first amplification in triplicate for each DNA sample. The PCRs were performed in 25-μL volumes with final concentrations of 1x supplied buffer (KAPA HiFi HotStart ReadyMix, Roche, Switzerland) and 0.3 μM of each forward and reverse primer (Supplementary Table 8). In total, 1 μL of extracted DNA was added, ranging in concentration from 2.35 to 143.98 ng/μL. A negative PCR control was carried out in triplicate, by adding 1 µL of PCR grade water instead of DNA sample, as well as positive PCR controls for 16S rRNA, 18S rRNA and rbcL, consisting of mock communities (Supplementary Table 5). The PCR program started with 95 °C for 3 min, followed by 18 cycles (16S rRNA), 29 cycles (*rbcL*) or 24 cycles (18S rRNA and rbcL) consisting of 95 °C for 20 s, 50 °C (16S rRNA), 58 °C (rbcL) or 56 °C (18S rRNA) for 15 s, and 72 °C for 15 s. A final extension of 72 °C for 5 min was performed and the PCR products from the three independent reactions for each sample were then pooled and cleaned. Each of the pooled reactions (a total of 75-μL) were cleaned using a 0.8x bead:sample volume ratio of selfmade SPRI beads and separated with a magnetic stand following the protocol of the Agencourt AMPure XP Kit (Beckman Coulter). The cleaned up PCR was stored at -20 °C until further processing.

The second PCR, consisting of a limited-cycle amplification, was carried out to add multiplexing indices and Illumina sequencing adapters. Each sample was dual-indexed by using the Nextera^®^ Index Kit A and D (Illumina, USA). The index PCR was performed in a 20-μL volume with final concentrations of 1x supplied buffer (KAPA HiFi HotStart ReadyMix, Roche, Switzerland) and 0.3 μM of each Nextera Index forward and reverse primer. Two mL of cleaned amplicons from the previous step were added. The PCR program started with 95 °C for 3 min, followed by 10 cycles consisting in 95 °C for 30 s, 55 °C for 30 s, and 72 °C for 30 s. A final extension of 72 °C for 5 min was performed and the PCR products were cleaned as described above.

The DNA concentration of the cleaned and indexed libraries was then determined using a Qubit (1.0) fluorimeter following recommended protocols for the dsDNA HS Assay. The libraries were then normalized and pooled at a 3.2 nM concentration. The pooled libraries were cleaned up twice and DNA concentration determined using a Qubit (1.0) fluorimeter. Absence of adapters was checked with a HS 1000 chip on a TapeStation device (Agilent). PHiX control was added at a 1% concentration. Paired end (2 × 300 nt) sequencing was performed on an Illumina MiSeq (MiSeq Reagent kit v3, 300 cycles) at the Genomic Diversity Centre (GDC) at the ETH, Zurich, Switzerland following the manufacture's run protocols (Illumina, Inc.). The MiSeq Control Software Version 2.2 including MiSeq Reporter 2.2 was used for the primary analysis and the de-multiplexing of the raw reads.

*Script for the ImageJ using custom-written macro and “Huang dark” auto-thresholding*

//run("Brightness/Contrast...");

setMinAndMax(145, 25000);

setAutoThreshold("Huang dark");

//run("Threshold...");

setThreshold(3500, 65535);

run("Analyze Particles...", " show=[Count Masks] display summarize");

#### **Supplementary Discussion**

#### Creation and validation of a specific reference database for single-species detection and quantification in the synthetic periphyton

Because the ability to accurately assign species in a sample is directly related to the quality of the reference database. We first created our own database for the twenty-six single species that were used to establish the synthetic periphyton and their respective amplicon gene sequences (Supplementary Table 6). Briefly, DNA was extracted from single species and genes of interest (18S rRNA, 16S rRNA and *rbcL* genes) were amplified in a PCR reaction (the list of primers is shown in Supplementary Table 8). Then the PCR products were purified and Sanger sequenced (Supplementary Table 6) (see Methods for more details). Obtained database of single species amplicon sequences was used to assign Amplicon Sequence Variants (ASVs) to single species. We tested the accuracy of the approach for single species identification and the quantification of their relative abundance by using a mock community composed of a defined mixture of extracted DNA from the twenty-six phototrophic species (Supplementary Table 5).

Sequencing of the 18S rRNA gene gave good resolution at the genus level, allowing detection of all green algae and all diatoms within the mock community (Supplementary Fig. 5A). However, by using these 18S rRNA primers it was not possible to distinguish two diatom species (*Fragilaria crotonensis* and *Fragilaria capucina*) and two green algae species (*Pediastrum duplex* and *Pediastrum boryanum*) of the same genus. We overcame the limitation for *Fragilaria* species by additionally sequencing *rbcL* gene. Note that depending on the used primers for amplicon sequencing, slightly different species abundances were observed in the mock community (Supplementary Fig. 5A and 5B). Although equal DNA amounts from all single species, except for *Cyclotella*, were added into the mock community (Supplementary Table 5), no equal abundances of these species were measured (Supplementary Fig. 5A and 5B). Moreover, not a double amount of *Cyclotella* was measured compared to the other species in the mock community. Together our data point at variations in gene copy numbers and/or differences in PCR primer efficiencies among the species [4]. With 18S rRNA gene primers the lowest abundance was detected for *Tabellaria* sp. (Supplementary Fig. 5A). Godhe *et al.* suggested that diatom cell length and cell biovolume could be a proxy for 18S gene copy number per cell due to a correlation between the cell size and gene copy numbers [4]. However, since the cell biovolume of *Tabellaria* is not the smallest from all species in our community (data not shown), the low abundance of *Tabellaria* probably results from poor PCR amplification efficiency from the DNA of this species rather than cell size.

Plastid large subunit of ribulose-1,5-bisphosphate carboxylase (rbcL) is a preferred locus in amplicon sequencing of diatoms [5], since it enables higher taxonomic resolution at the species level and discrimination between species of the same genus. When we used *rbcL* barcode, only diatom species were detected in the mock community, confirming the specificity of the primers (Supplementary Fig. 13). By using chloroplast-encoded *rbcL* gene, we also measured the highest abundance of *Cyclotella meneghiniana*. This was expected, since double DNA amount of this species was added to the mock community and also *Cyclotella* *meneghiniana* harbors high chloroplast numbers and thus many *rbcL* genes. However, also with *rbcL* gene barcode, we did not measure equal abundances among the other species in the mock community although equal DNA amounts were used. The lowest abundances were detected for *Nitschia* and *Sellaphora* genera, which could be explained by their small cell biovolume and low chloroplast numbers per cell [6]. Indeed, Vasselon *et al.* showed a correlation between the number of copies of the *rbc*L gene per cell and the diatom biovolume [6]. Our results fit this correlation with the smallest cells (e.g., *Nitzschia* and *Sellaphora)* and largest cells (e.g., *Cymbella*, *Fragilaria* and *Gomphonema)* showing the lowest and highest abundances, respectively (Supplementary Fig. 5B).

Among the four cyanobacteria species that we tested, double DNA amount of *Pseudanabaena galeta* was added into the mock community, in order to check the ability to obtain respective community composition with the selected 16S rRNA primers [1]. Our results show that by using 16S rRNA gene as a barcode, approximately double amount of this species compared to other cyanobacteria was measured (Supplementary. Fig. 5C). On the other hand, although equal DNA amount of *Chamaesiphon polonicus*, *Merismopedia glauca* and *Phormidium sp.* was added to the mock community, their relative abundances differed, suggesting variability in 16S rRNA gene copy numbers [7]. Note that we also detected amplification from the chloroplast DNA, which reflects non-specific binding of the used 16S rRNA primers.

In summary, our results show that 18S rRNA gene primers provide resolution sufficient for green algae and diatoms at the genus level, whereas *rbcL* primers enabled higher taxonomic resolution for diatoms, down to the species level. Moreover, the 16S rRNA primers allowed distinguishing the four cyanobacteria species from each other, although an unspecific amplification from the chloroplast was also detected. Depending on the primers used for amplicon sequencing, slightly different single species abundances were observed in the mock community due to variations in gene copy numbers and/or primer specificities. This demonstrates the importance to test for the precision of the selected primers in the amplicon sequencing approach for species quantification before performing community composition analysis. It also shows that targeting more than one gene, even within one phototrophic group, assures accurate community characterization at the species level.

#### **Supplementary Figures**

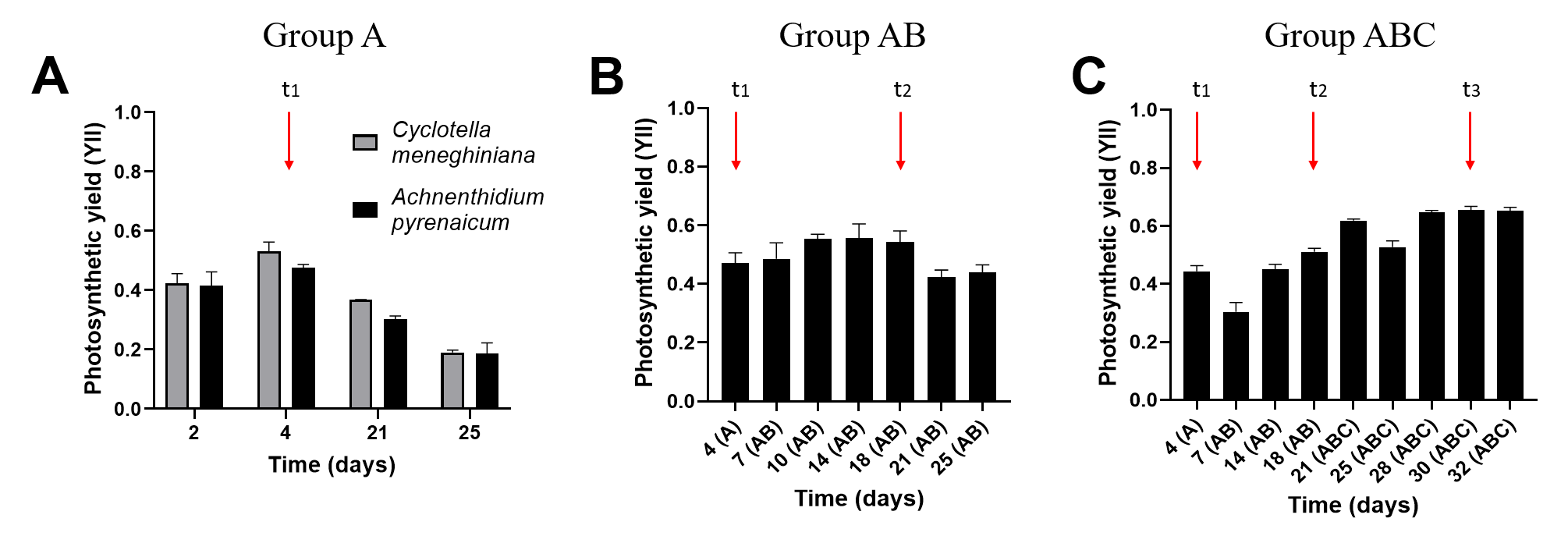

**Supplementary Fig. 2.** Measurement (mean ± standard deviation, n = 5) of the photosynthetic quantum yield of group A **(A)**, group AB **(B)** and group ABC **(C)** species. Photosynthetic quantum yield was measured by using iPAM (imaging Pulse-Amplitude-Modulation M-series) chlorophyll fluorimeter. Red vertical arrows indicate selected time points when group B species were added on top of group A (t_1_), group C on top of group AB (t_2_) and periphyton was established (t_3_).

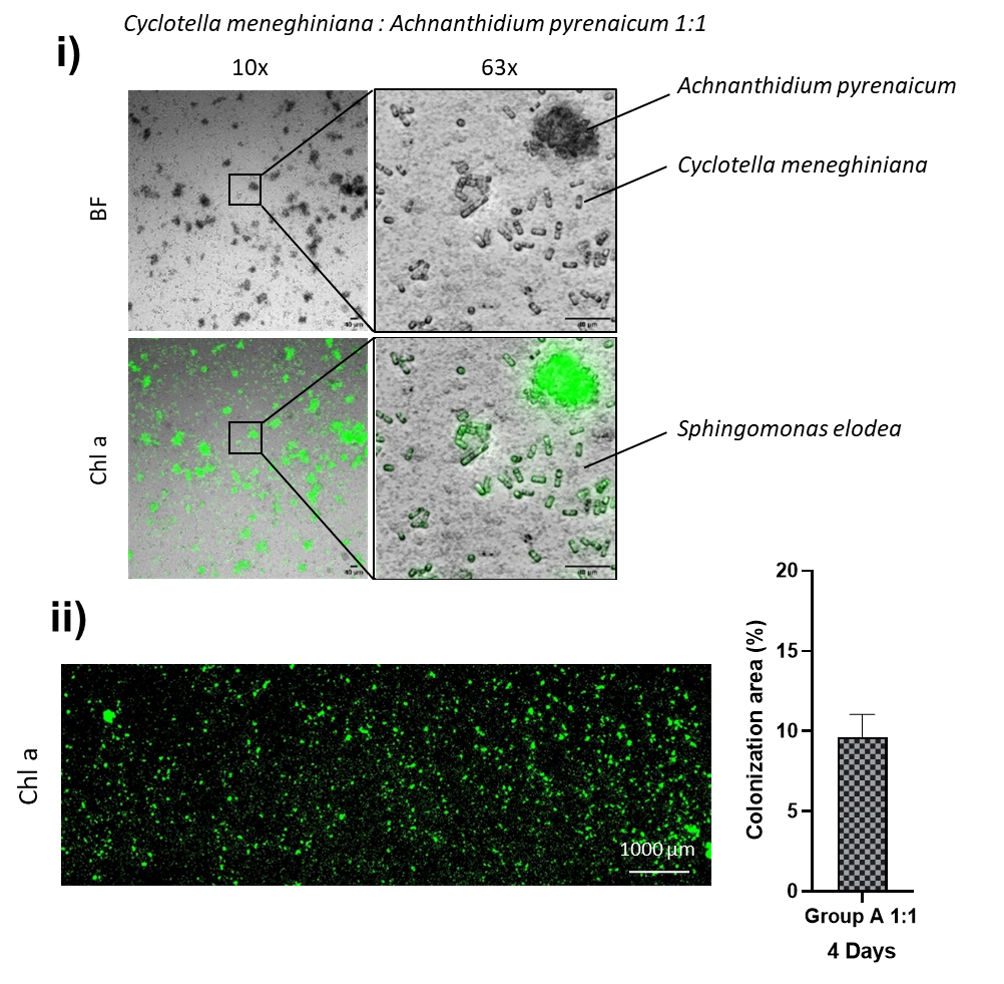

**Supplementary Fig. 3.** Colonization of group A diatom species *Cyclotella meneghiniana* and *Achnanthidium pyrenaicum* together at 1:1 ratio **(i)**. 9 x 3 microscopy images were acquired, montaged and stitched together in order to calculate the percentage of surface colonized area (mean ± standard deviation, n = 3) **(ii)** (see Methods for more details)*.*

**i)**

**ii)**

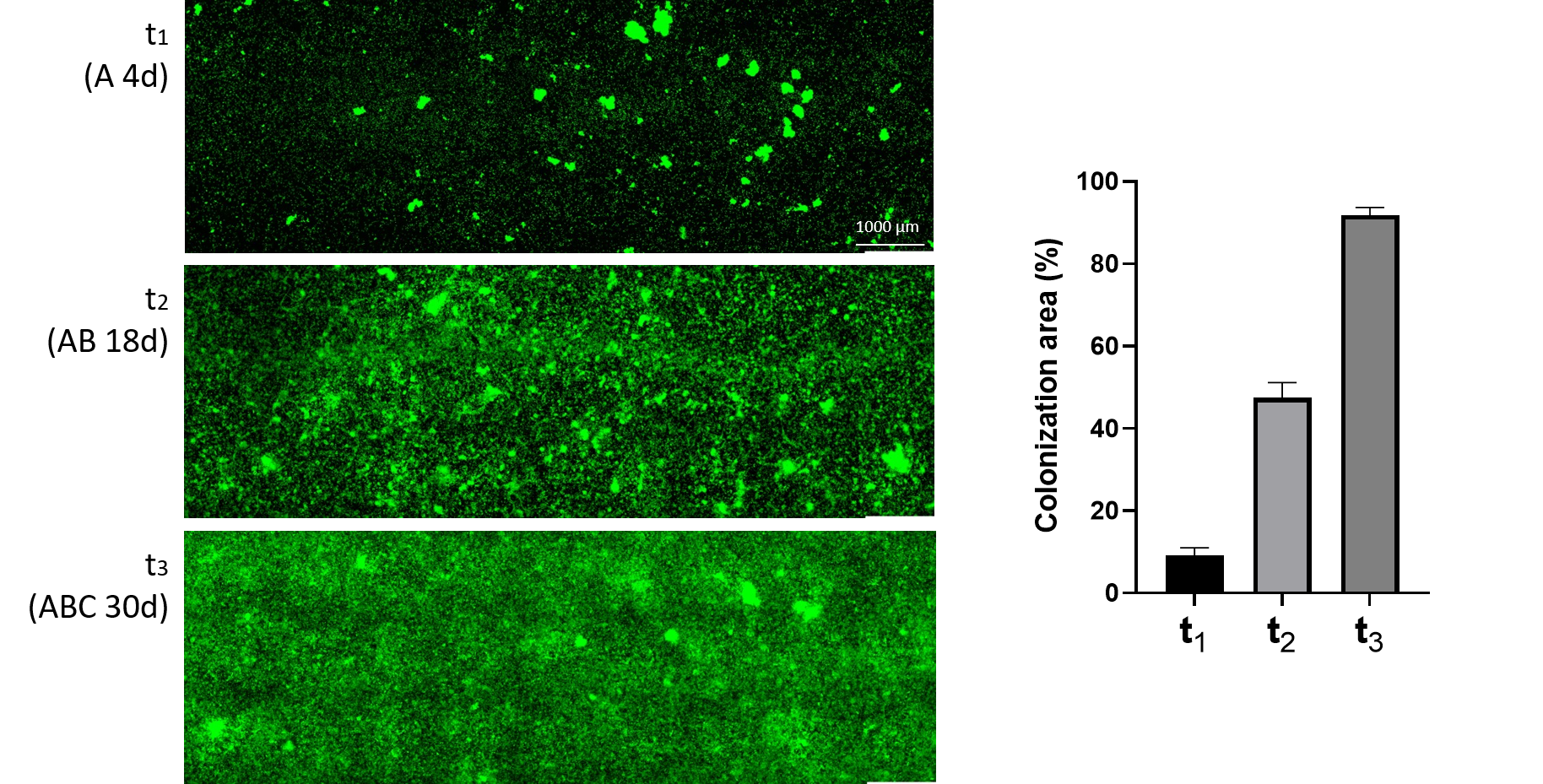

**Supplementary Fig. 4:** Example of representative microscopy images **(i)** showing surface colonization by the phototrophic species during periphyton growth at t_1_ (4 days), t_2_ (18 days) and t_3_ (30 days) (scale bar 1000 µm). **(ii)** Microscopy images (3 replicates from 3 independent experiments) were used to calculate the percentage of surface colonized area (mean ± standard deviation, n =9).

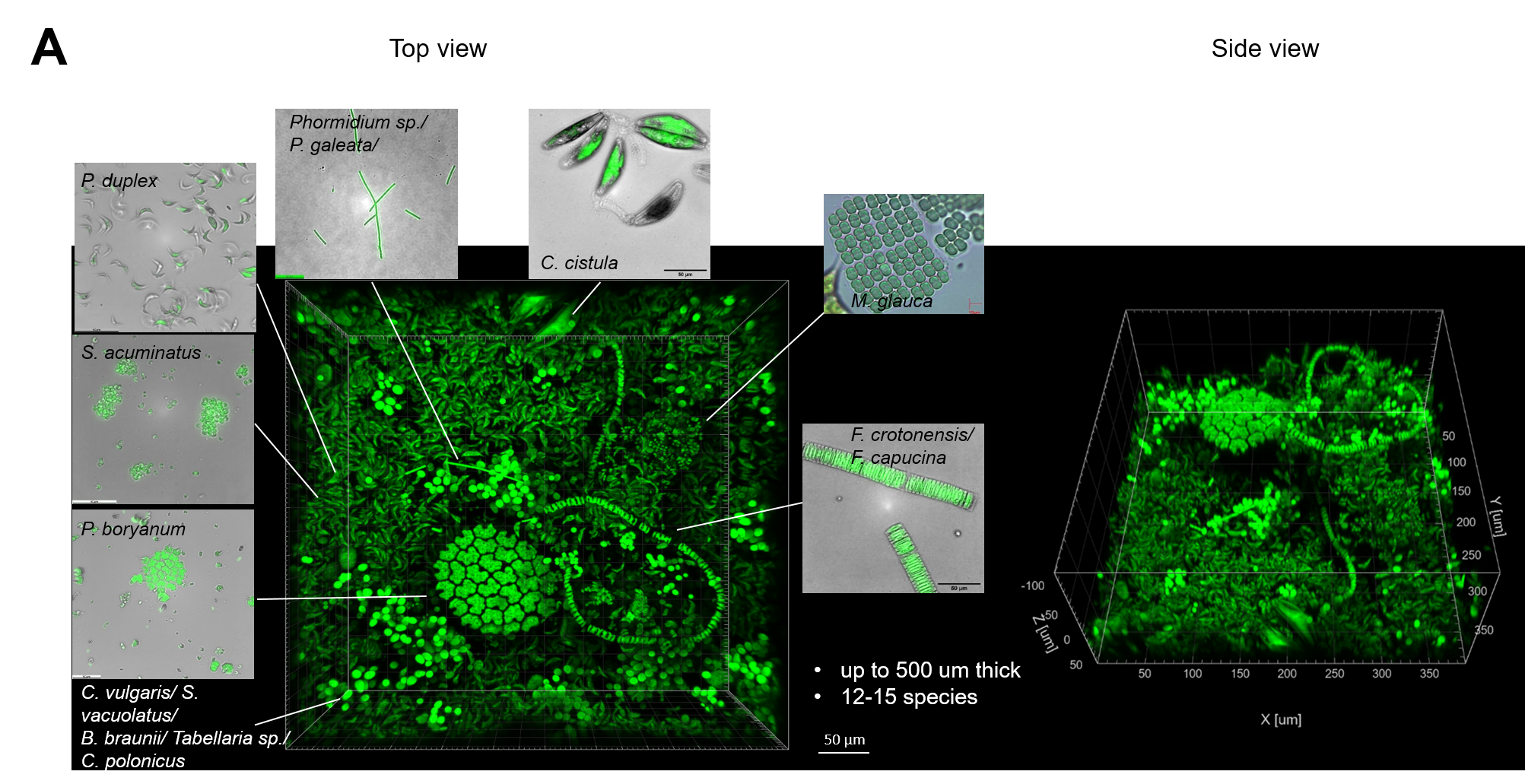

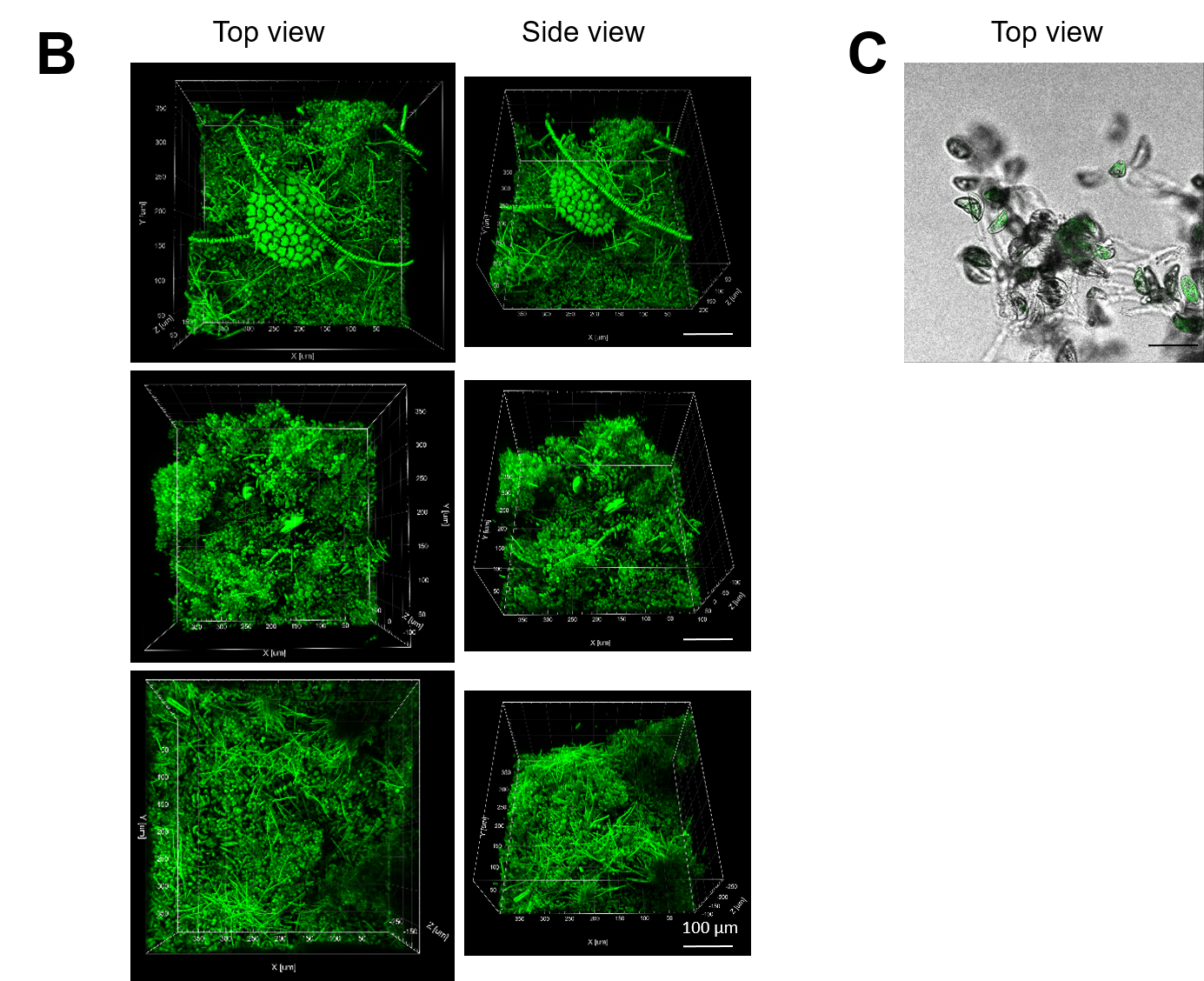

**Supplementary Fig. 5:** Representative confocal laser scanning microscopy images of the established periphyton community at 30 days (t_3_) with 12-15 identified phototrophic species based on their morphology (scale bar: 100 µm) **(A)**. Additional species (i.e., *C. cuspidata*, *C. accomoda*, *Melosira sp*., *N. vermicularis*, *Synedra sp*. and *Ulnaria ulna*) were also identified from other images (not shown). The green color indicates autofluorescence of chlorophyll *a* (excitation at 633 nm and emission at 650-750 nm). Microscopic observations were performed on at least five different points of one glass slide. Other representative images are shown in **(B)** (scale bar: 50 µm). Representative image showing stalks and extracellular polymeric substances produced by species of the community obtained with T-PMT brightfield channel **(C)**.

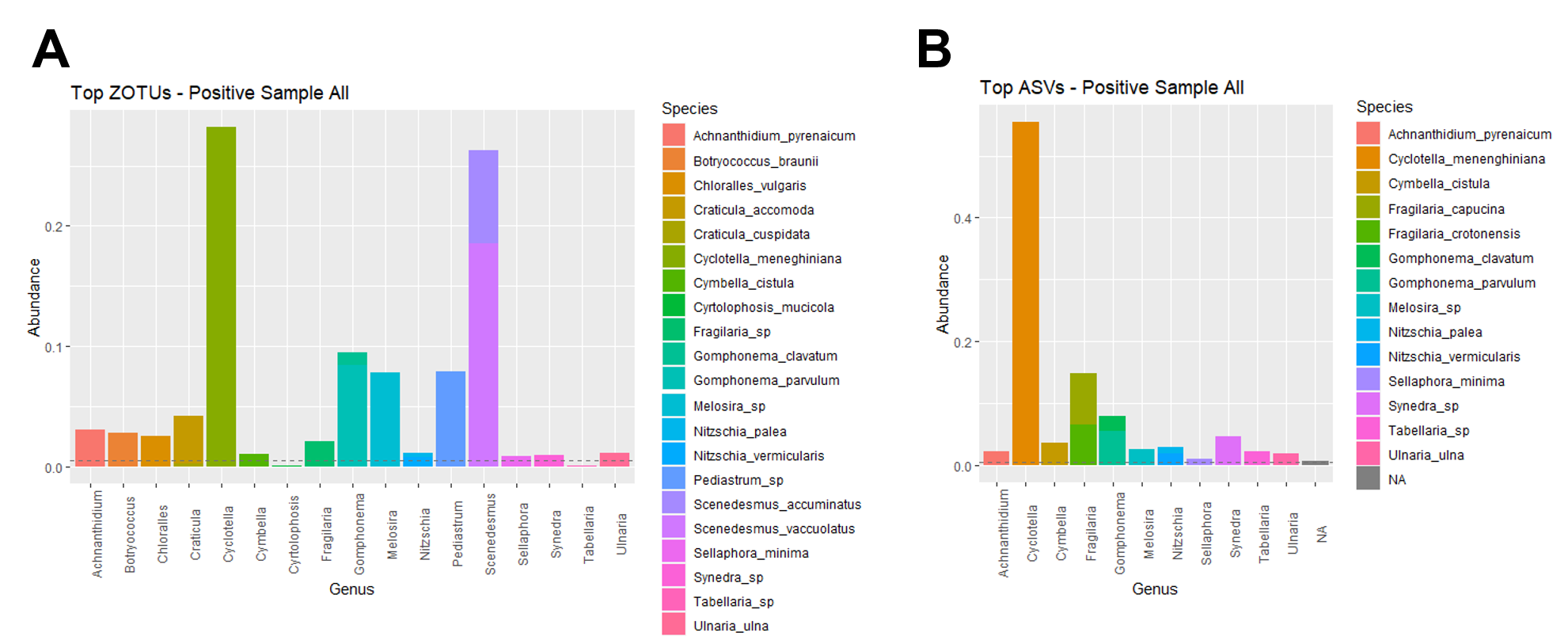

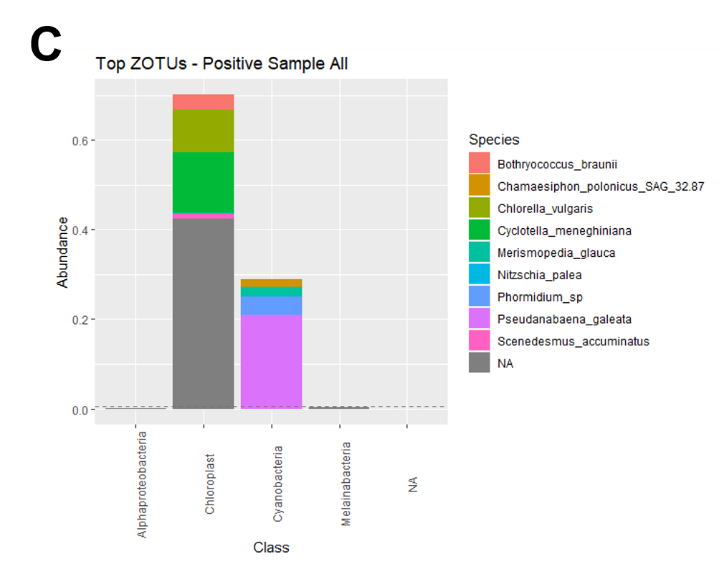

**Supplementary Fig. 6.** Abundances of the single phototrophic species in the mock community from taxonomy-based clustering at the genus or class level of assigned 18S rRNA **(A)**, *rbcl* **(B)** and 16S rRNA **(C)** gene sequences. 200 ng DNA from *Cylotella meneghiniana* and *Pseudanabaena galeata* each and 100 ng DNA from each of the 24 remaining single species were pooled. NA: not assigned species. Dashed horizontal line represents relative abundance of 0.01.

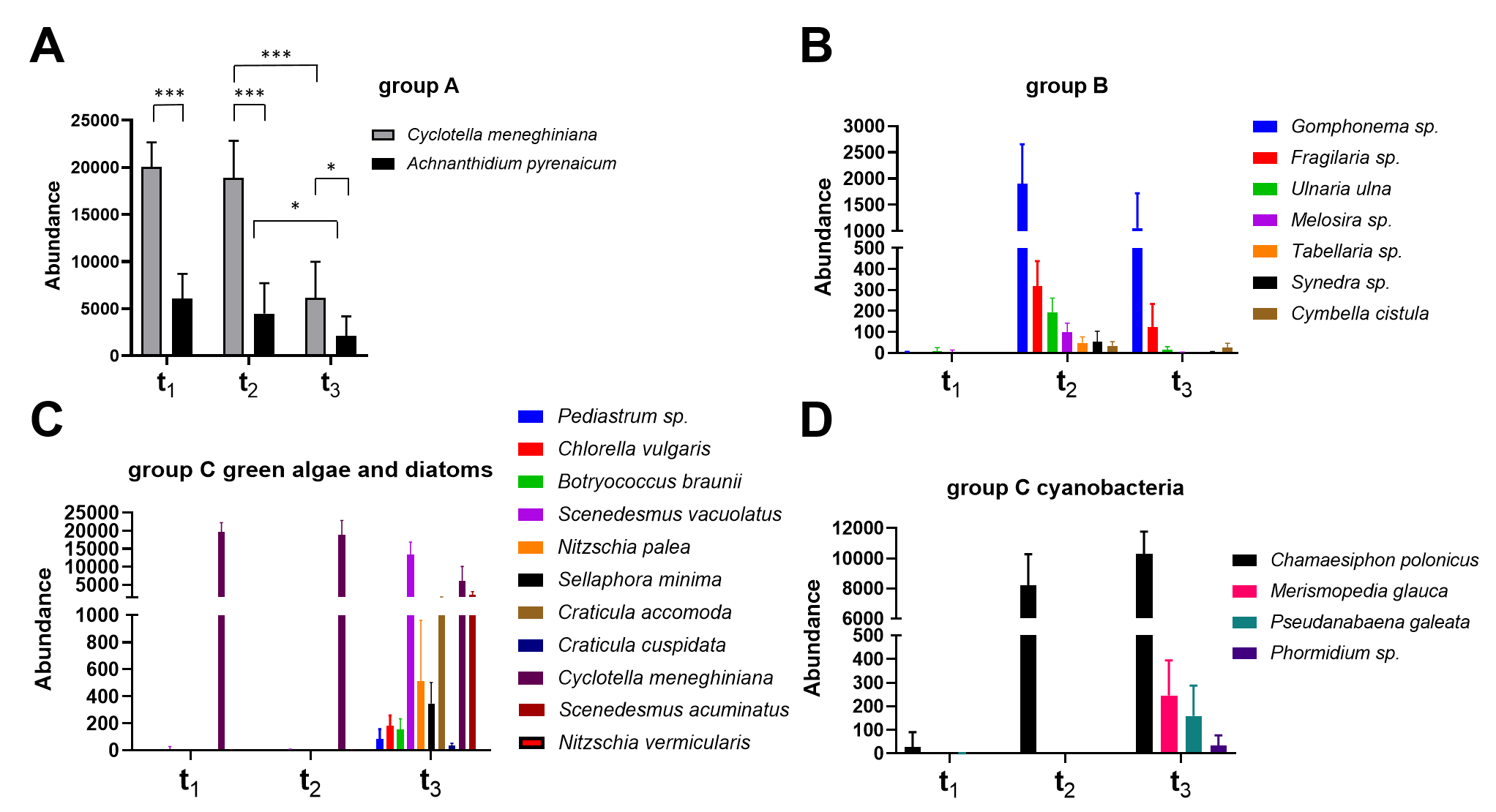

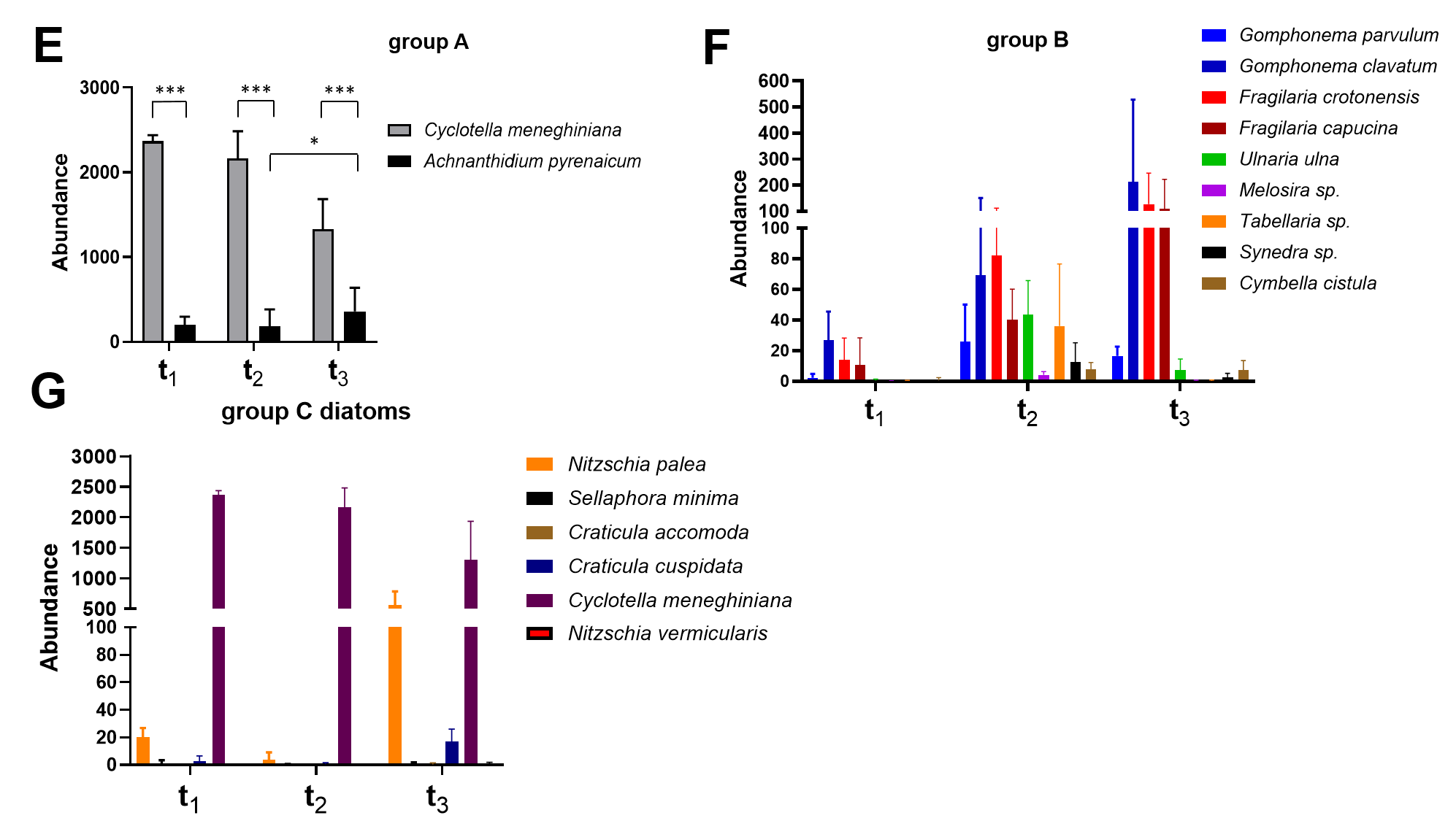

**Supplementary Fig. 7.** Synthetic periphyton composition during periphyton establishment at the species level measured with 18S rRNA (**A-C**), 16S rRNA (**D**) and *rbcL* gene sequencing (**E-G**). Shown is the community composition profile and abundances inferred from taxonomy-based clustering at species or genus level of assigned genes at 4 (t_1_), 18 days (t_2_) and 30 days (t_3_) during periphyton formation (mean ± standard deviation, n = 9). Abundances of group A **(A)**, group B **(B)** and group C **(C)** eukaryotic species (green algae and diatoms) measured using 18S rRNA amplicon sequence were measured at t_1_, t_2_ and t_3_ during periphyton establishment. Abundances of group C cyanobacteria **(D)** were measured using 16S rRNA. Finally, abundances of group A **(E)**, group B **(F)** and group C **(G)** diatom species were measured with *rbcL*. ns – not significant, * P<0.05, ** P<0.005 *** P<0.0001, one-way ANOVA test followed by Tukey’s test. Using *rbcL*, we detected more than two group A species at t_1_, pointing at non-specific binding of *rbcL* primers. However, due to low concentrations of extracted DNA from timepoint t_1_ and since the overall abundance of non-specific counts was less than 5% of the community, these counts were treated as noise.

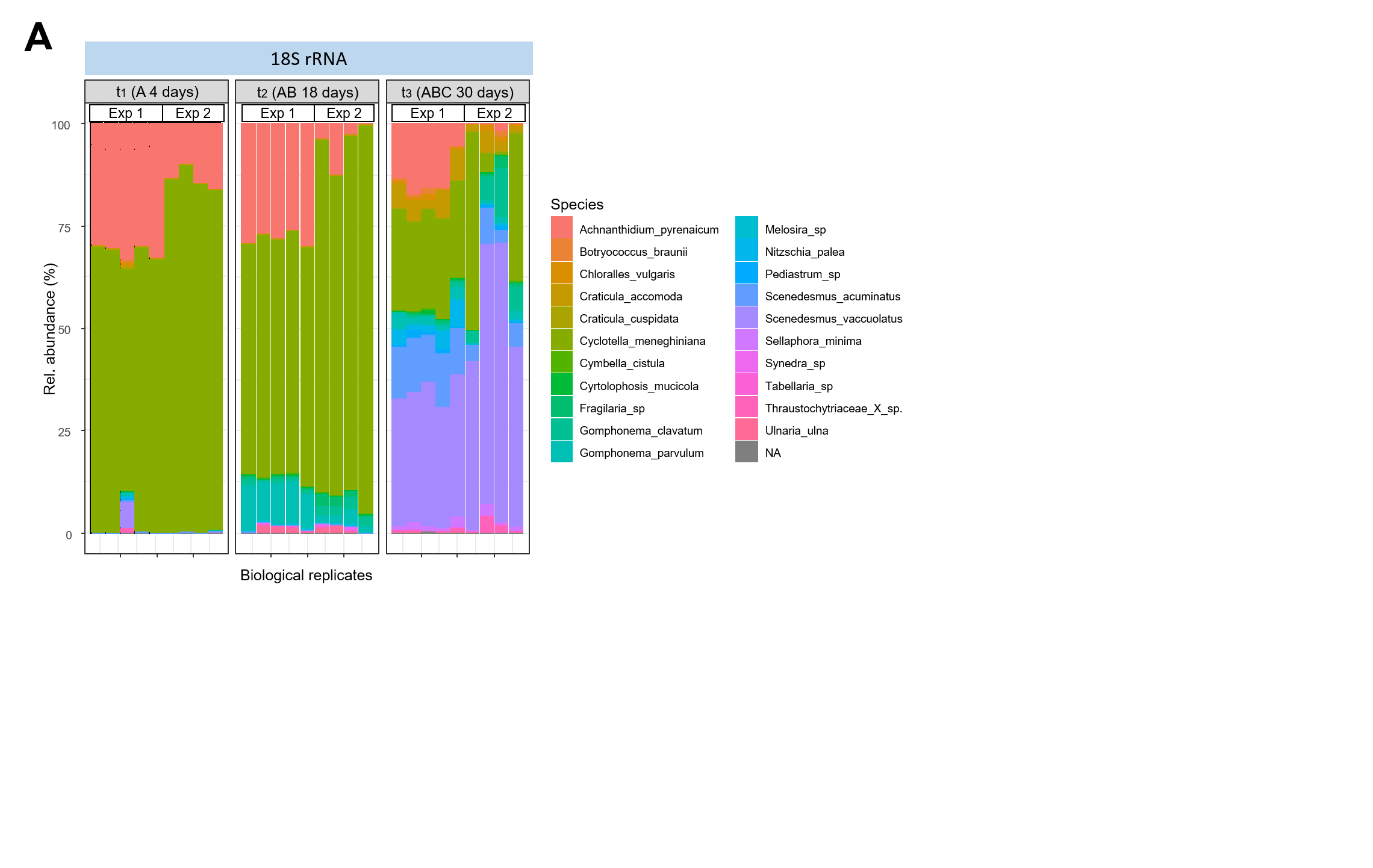

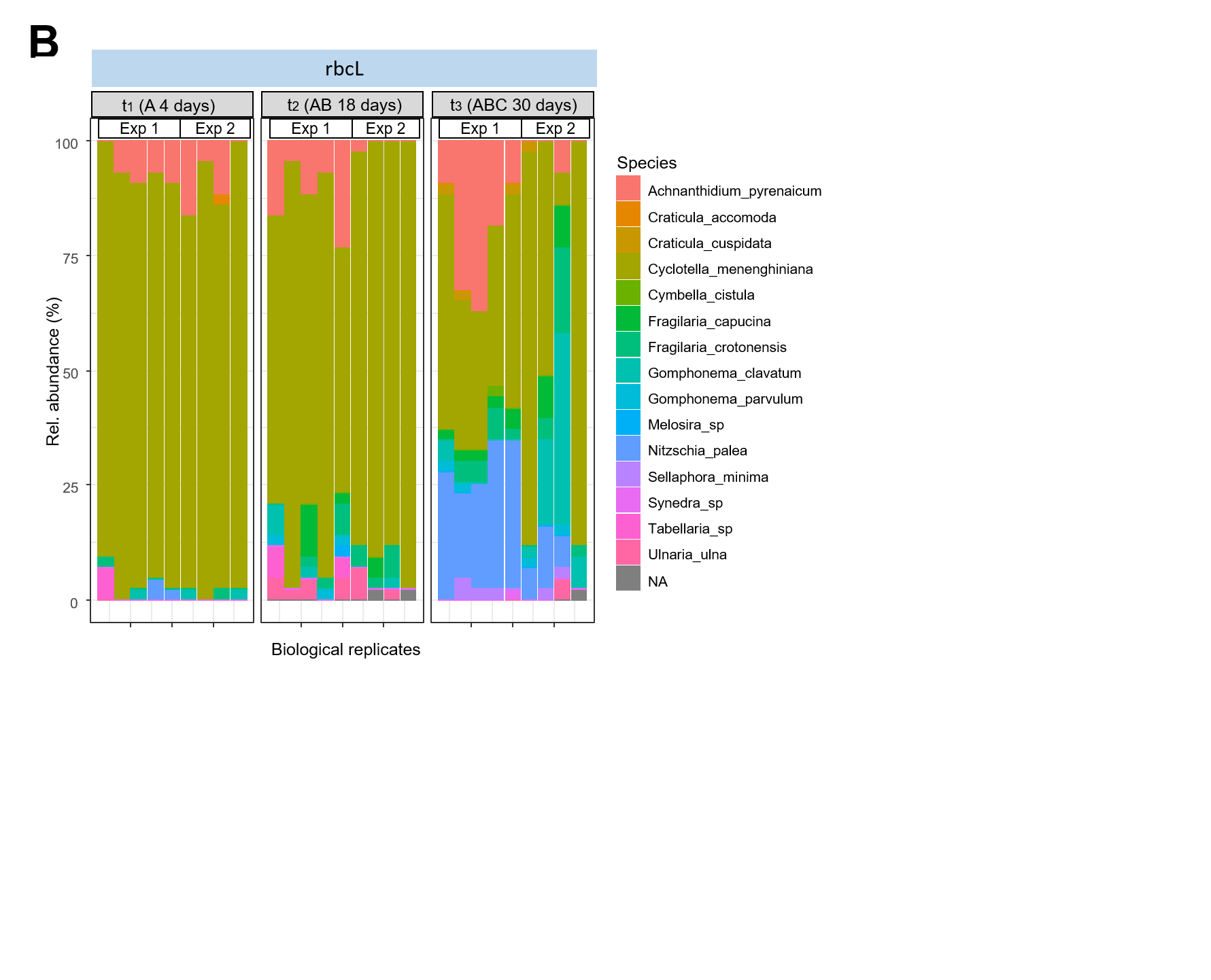

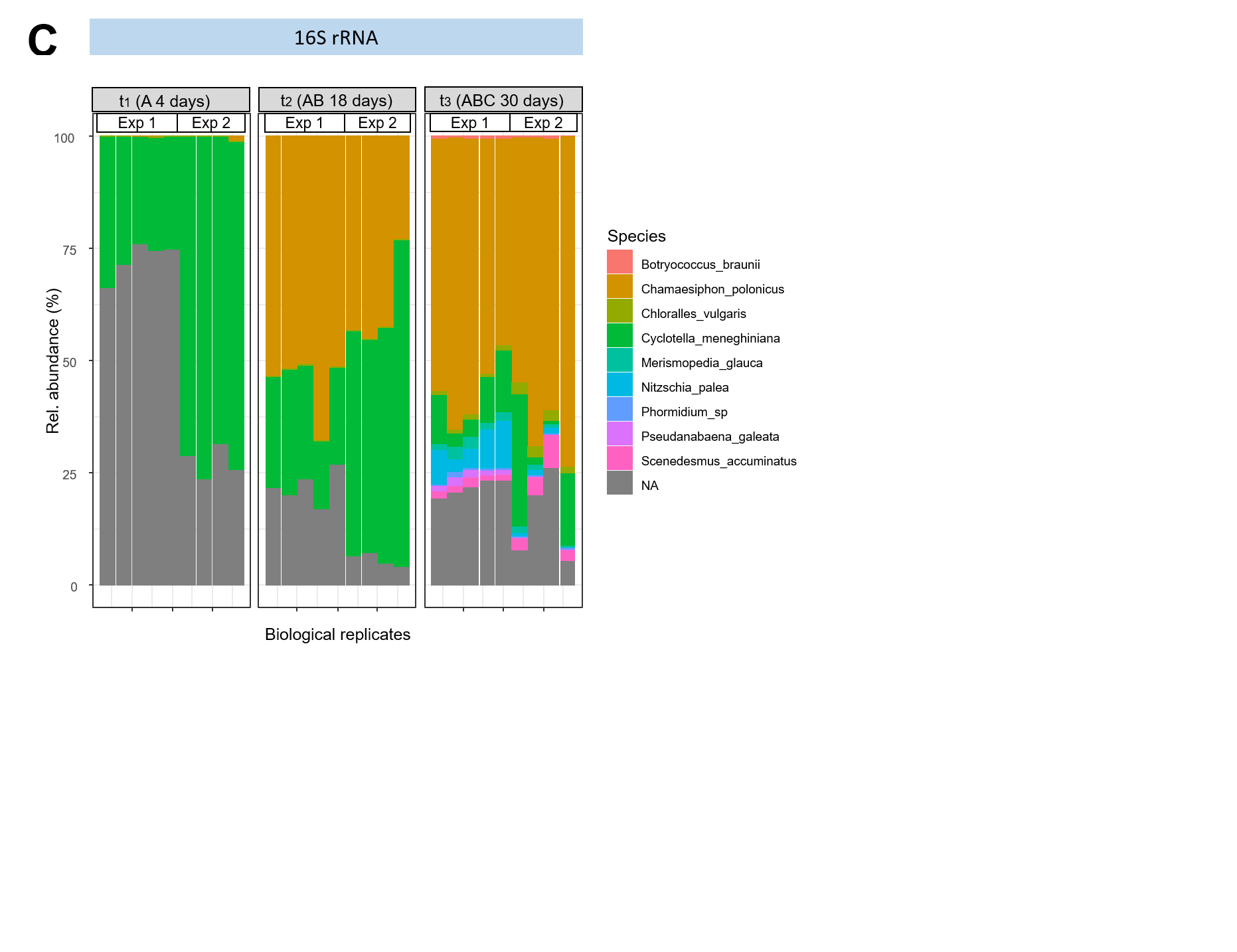

**Supplementary Fig. 8.** Reproducibility of the community composition during periphyton establishment. Data are obtained from two independent experiments (Exp 1 and Exp 2, with n = 5 and 4 biological replicates, respectively). Profile and relative abundances were inferred from taxonomy-based clustering at species level of assigned 18S rRNA **(A)**, *rbcl* **(B)** and 16S rRNA **(C)** genes at 4 (t_1_), 18 (t_2_) and 30 days (t_3_) during periphyton formation. NA: not assigned species.

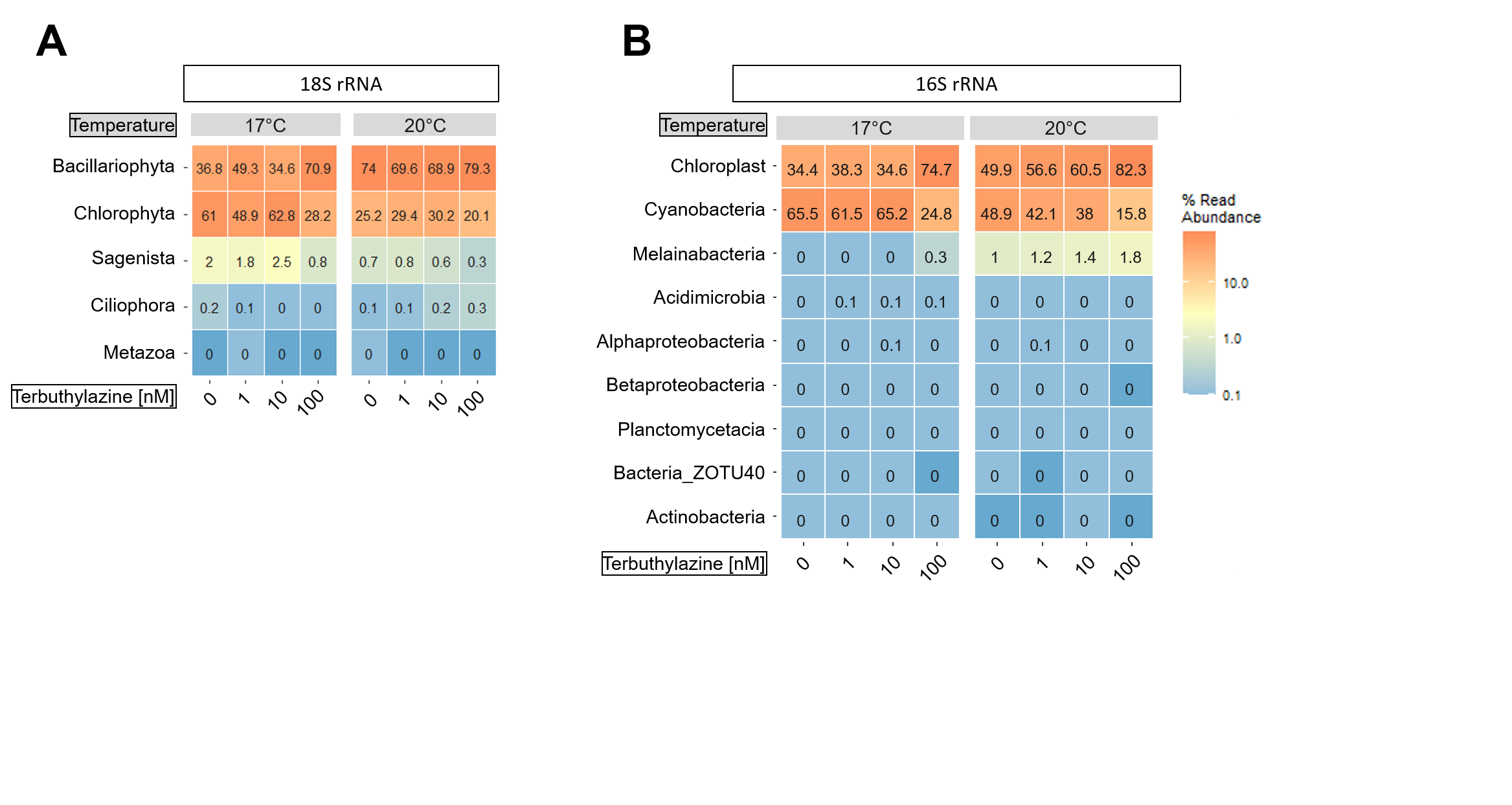

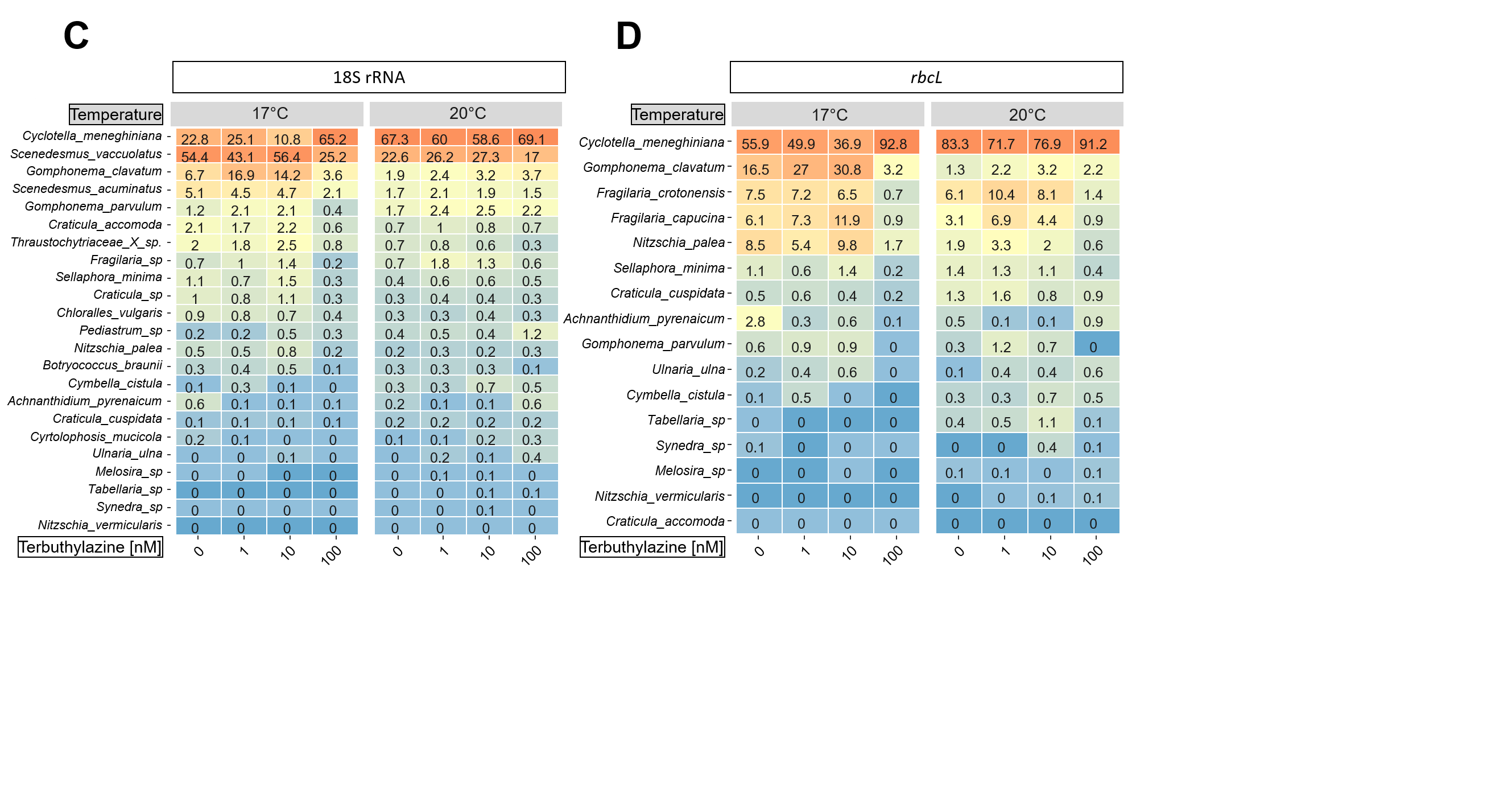

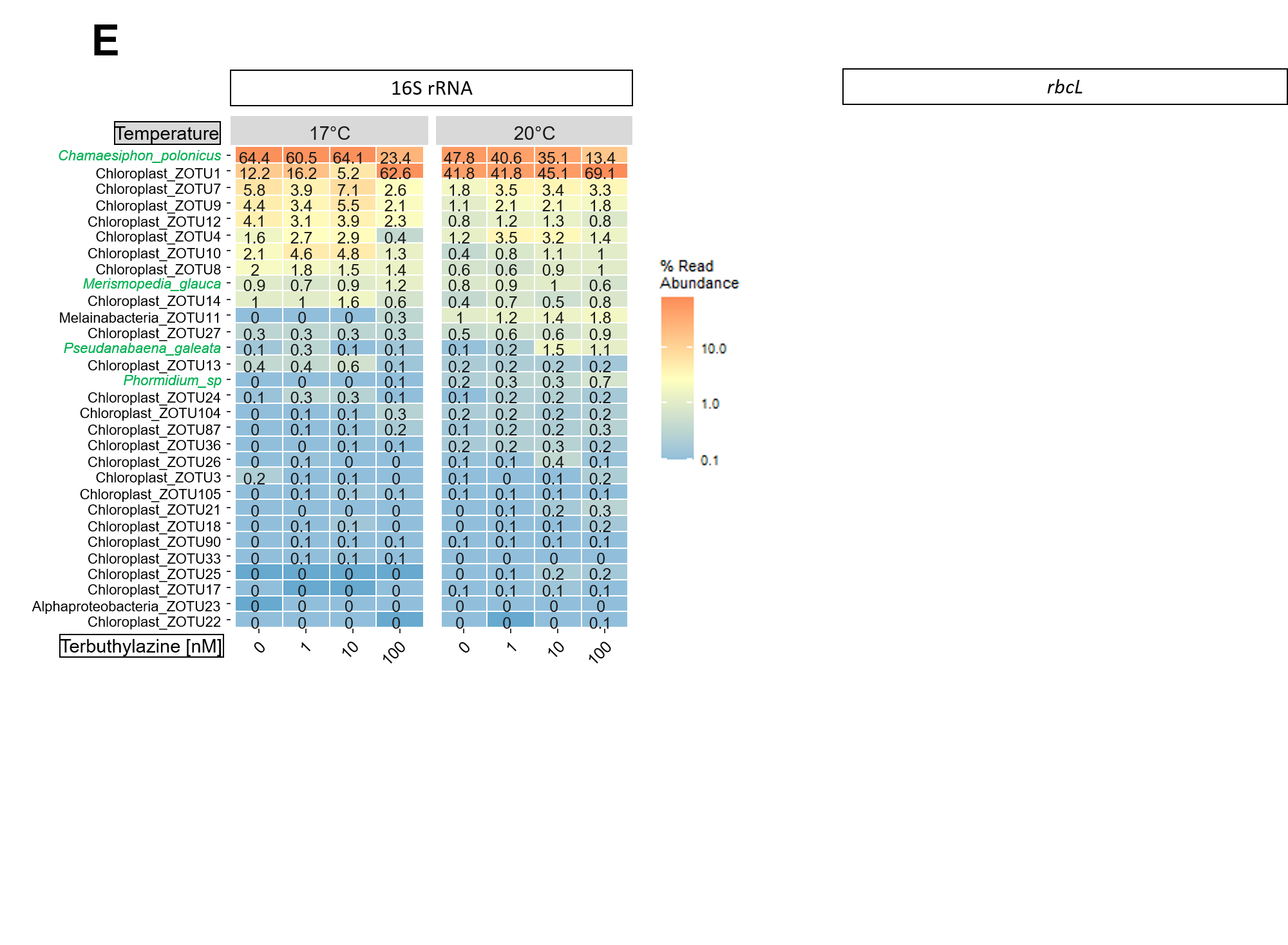

**Supplementary Fig. 9. Heatmaps showing relative abundances of the phototrophic species in established periphyton under different conditions.** Effects of terbuthylazine (0, 1, 10 or 100 nM) and temperature (17 °C or 20 °C) are shown at t_3_ (ABC 30 days) at the phylum level measured with 18S rRNA **(A)** and 16S rRNA **(B)** and at the species level measured with 18S rRNA **(C)**, *rbcL* **(D)** and 16S rRNA (**E**) genes. Species labelled in green correspond to the 4 cyanobacteria that we used to establish the periphyton. Data are based on the average of Amplicon Sequence Variants (ASVs) abundance (n = 3-4).

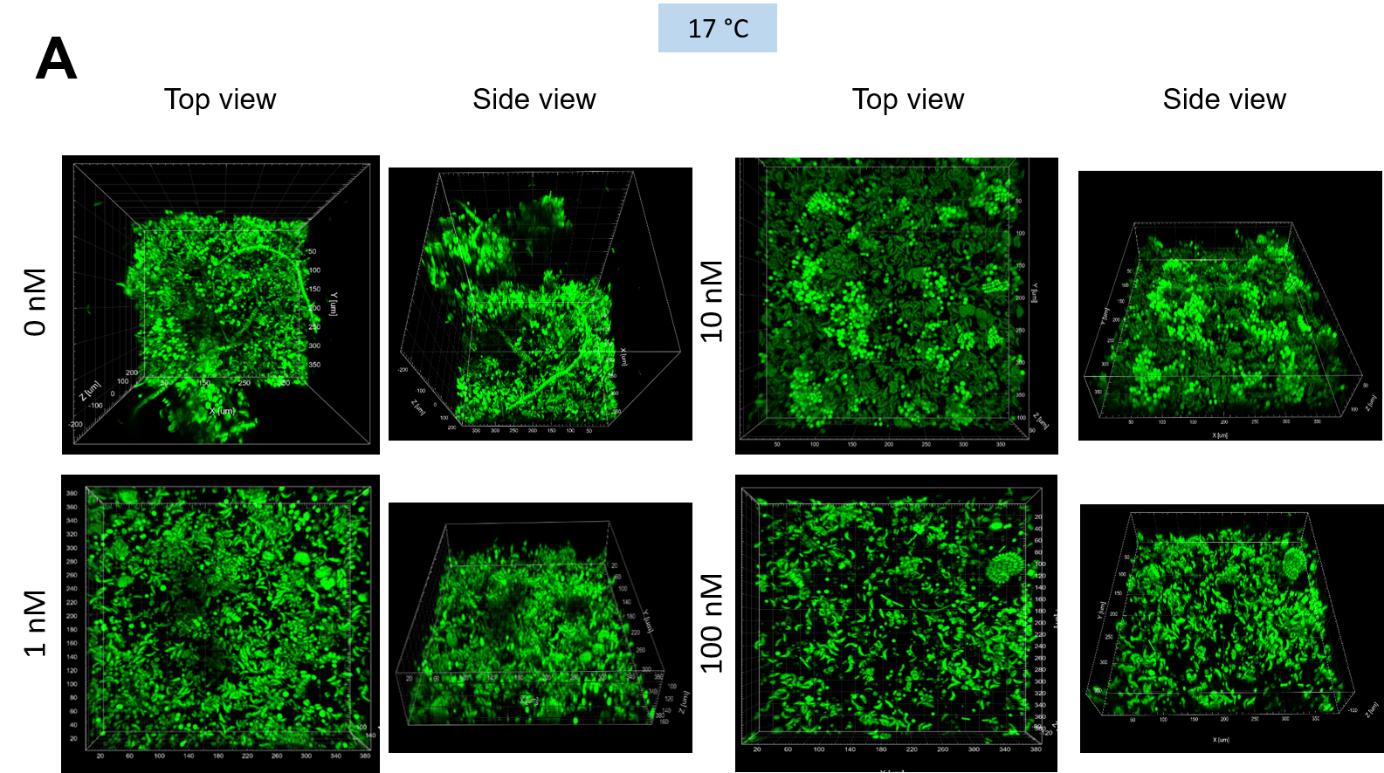

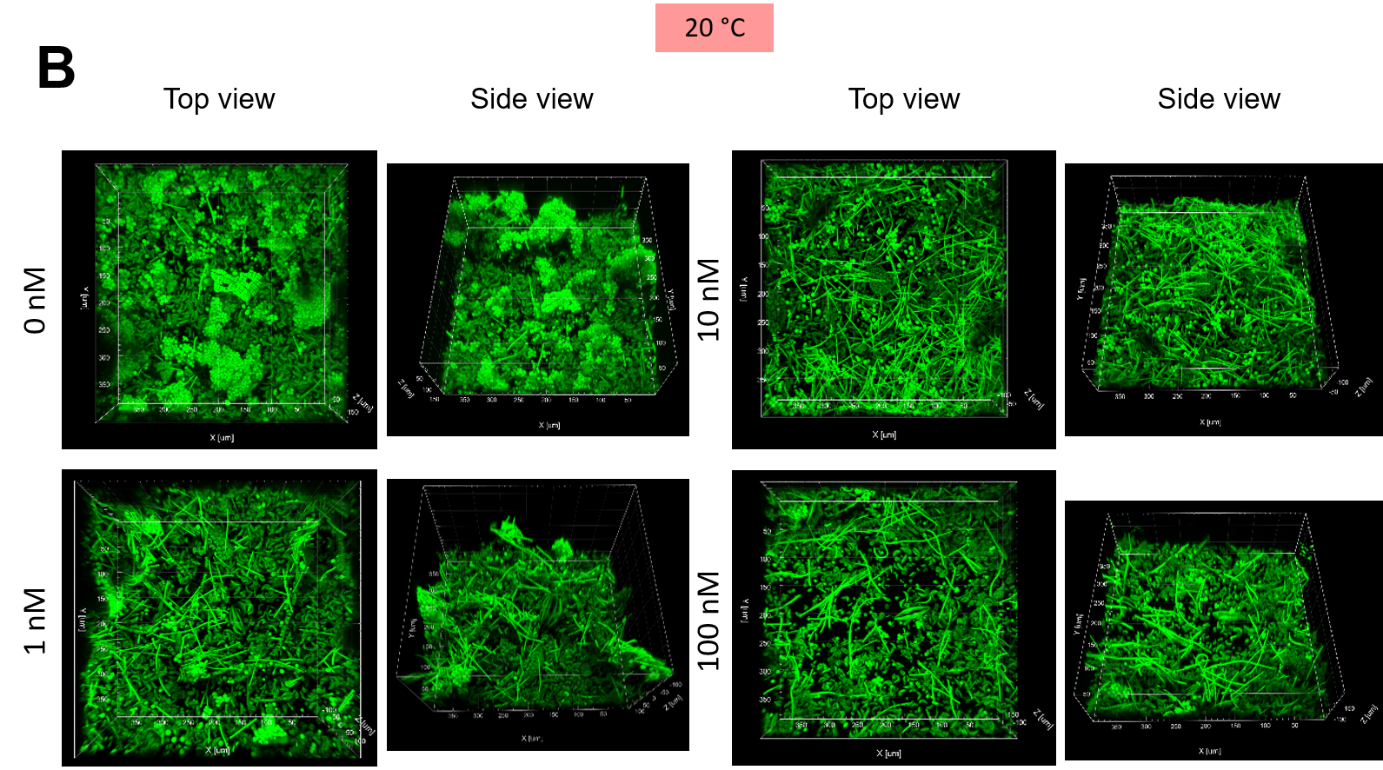

**Supplementary Fig. 10:** Representative confocal laser scanning microscopy images of established periphyton community at 30 days (t_3_) under different treatments. The treatments correspond to 4 levels of the herbicide terbuthylazine (0 1, 10 and 100 nM) and 2 different temperatures (17 and 20 °C).

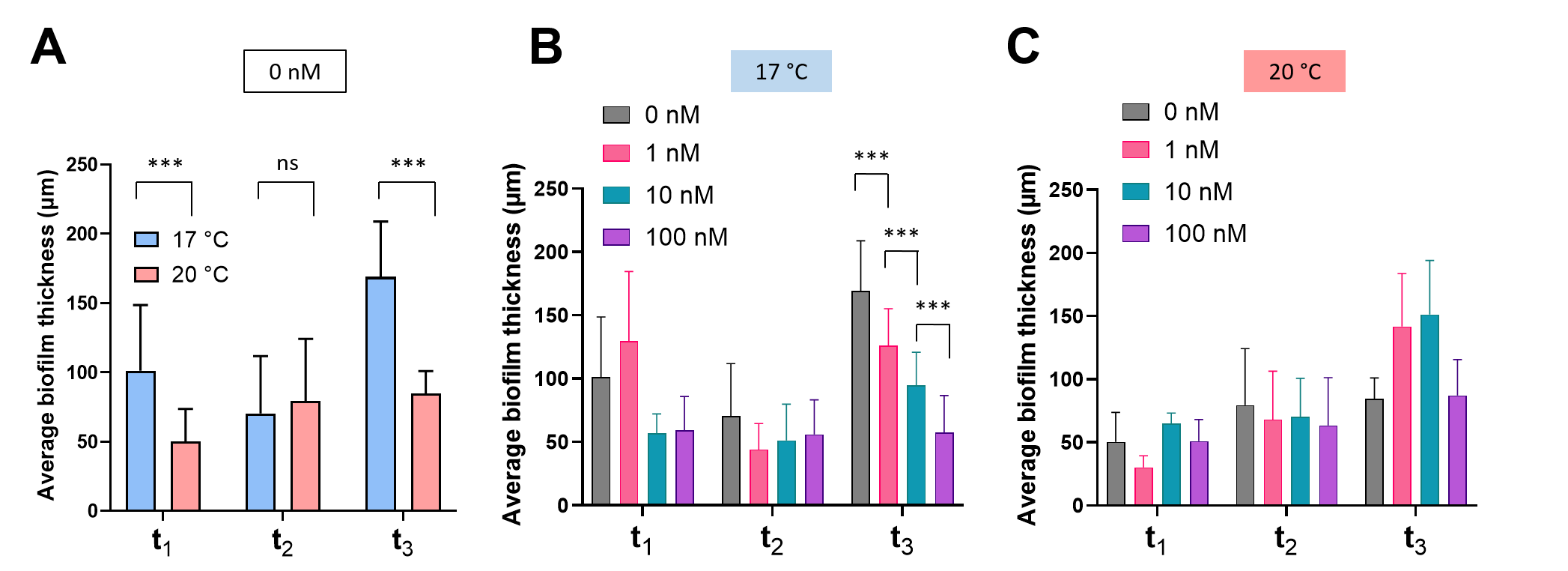

**Supplementary Fig. 11:** Average biofilm thickness of periphyton grown under different conditions. OCT measurements were obtained over 30 days of periphyton growth at time points t_1_ (A 4 days), t_2_ (AB 18 days) and t_3_ (ABC 30 days). Average biofilm thickness of periphyton grown under no terbuthylazine exposure at 17°C or 20°C is shown in **(A)**. The conditions correspond to periphyton grown in the presence of 0 nM (control), 1 nM, 10 nM or 100 nM terbuthylazine and at 17 °C **(B)** or 20 °C **(C)**. All experiments were performed in triplicate (mean ± standard deviation, n = 3), average biofilm thickness was determined from 20-30 images. ns – not significant, * P<0.05, ** P<0.005 *** P<0.0001, based on Students t-test for (A) and post hoc Tukey’s test for (B) and (C).

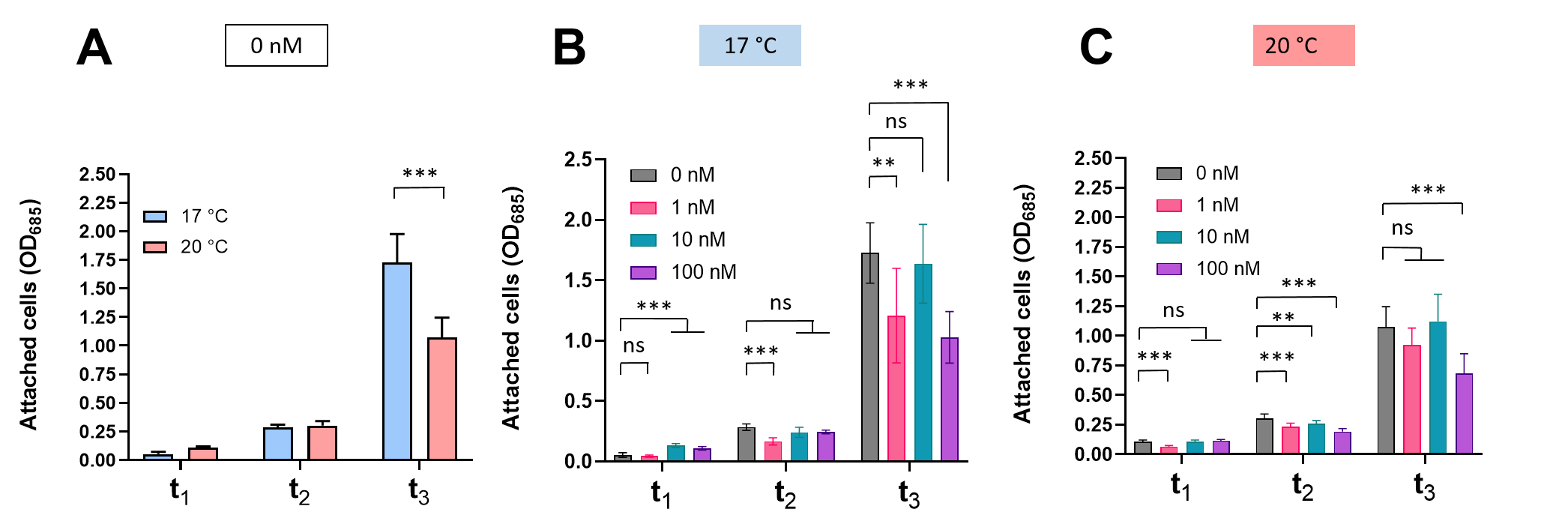

**Supplementary Fig. 12:** Benthic biomass of periphyton grown under different conditions. Attached cells were measured with the spectrophotometer at 685 nm over 30 days of periphyton growth at time points t_1_ (A 4 days), t_2_ (AB 18 days) and t_3_ (ABC 30 days). Benthic biomass of periphyton grown under no terbuthylazine exposure at 17°C (blue) or 20°C (red) is shown in **(A)**. Benthic biomass of periphyton grown in the presence of 0 nM (control), 1 nM, 10 nM or 100 nM terbuthylazine and at 17 °C **(B)** or 20 °C **(C)** are also shown. All experiments were performed in triplicate (mean ± standard deviation, n = 3). ns – not significant, * P<0.05, ** P<0.005 *** P<0.0001, based on Students t-test for (A) and post hoc Tukey’s test for (B) and (C).

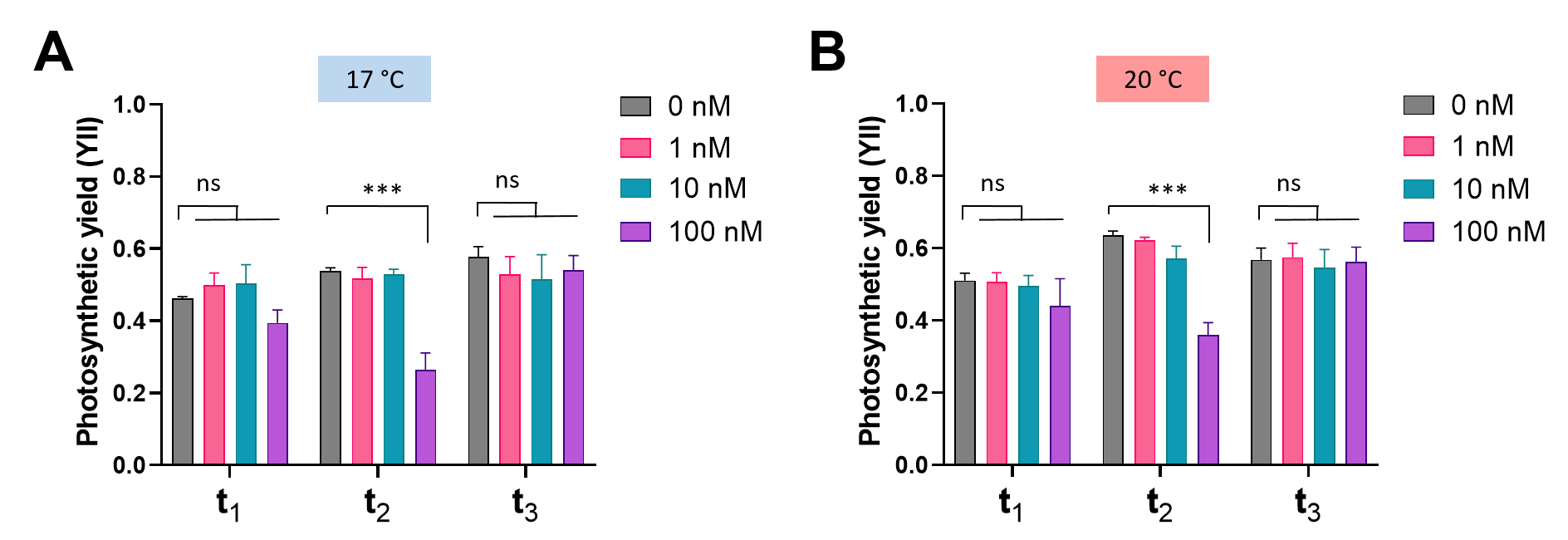

**Supplementary Fig. 13.** Photosynthetic yield of periphyton grown under different conditions. Photosynthetic efficiency measurements were obtained over 30 days of periphyton growth at time points t_1_ (A 4 days), t_2_ (AB 18 days) and t_3_ (ABC 30 days). The conditions correspond to periphyton grown in the presence of 0 nM (control), 1 nM, 10 nM or 100 nM terbuthylazine and at 17 °C **(A)** or 20 °C **(B)**. Photosynthetic yield (ΔF/F_m_′) was measured as described in 2.3.3. Material and Methods. All experiments were performed in 5 replicates (mean ± standard deviation, n = 5). ns – not significant, * P<0.05, ** P<0.005 *** P<0.0001, based post hoc Tukey’s test.

### **Supplementary tables**

See “Lamprecht_SI_Tables.xlsx” file.
